# Structural mechanisms for the recruitment of factor H by *Streptococcus pyogenes*

**DOI:** 10.1101/2025.08.05.668778

**Authors:** Amit Kumar, Kuei-Chen Wang, Partho Ghosh

**Affiliations:** Department of Chemistry and Biochemistry, University of California, San Diego, La Jolla, CA, 92093 USA; Biolegend, 8999 Biolegend Way, San Diego, CA 92121 USA

**Keywords:** *Streptococcus pyogenes*, M protein, FbaA, Factor H

## Abstract

The major bacterial pathogen *Streptococcus pyogenes* (Group A *Streptococcus,* or Strep A) recruits the negative regulator of the alternative complement pathway factor H (FH) to its surface. Antigenically sequence variable regions of several Strep A M proteins, including M5 and M6, bind FH but have no obvious sequence homology. A second Strep A surface-localized protein, FbaA, binds FH through a purported coiled-coil region, suggesting mimicry of the well-known coiled coil of M proteins. We determined the structures of fragments of M5 protein, M6 protein, and FbaA complexed with FH domains 6 and 7 (FH(6–7)). M5 and M6 proteins formed dimeric α-helical coiled coils, as expected, while FbaA instead consisted of a monomeric three-helix bundle preceded by a loop. FH(6–7) accommodated different binding modes in these three proteins, with very few common interacting amino acids. Based on contributions to binding, distinct FH-binding sequence patterns were constructed for M5 and M6 proteins, enabling identification of FH-binding sequences in M or M-like Enn proteins in 32 strains of differing M types. While FbaA was allelically sequence variable, its critical FH-binding amino acids were absolutely conserved in 95 strains of differing M types. Together, FH-binding sequences were identified in about half of the known 250 Strep A strains, with the majority due to FbaA. Our structural and functional elucidation of the mechanism of FH recruitment is applicable to precise investigation of its role in Strep A virulence.

## Introduction

*Streptococcus pyogenes* (Group A *Streptococcus,* Strep A) is responsible for morbidity and mortality on a global scale (1). This gram-positive bacterial pathogen is proficient at evading the human immune response, including the immediate response of the complement system, which crucially involves deposition of the opsonin C3b on the Strep A surface. Deposition leads to phagocytic uptake and destruction of the bacterium by neutrophils and macrophages. Strep A recruits human proteins to its surface to block C3b deposition (2). Among these are the blood clotting protein fibrinogen (3, 4) as well as the negative regulator of the classical and lectin pathways of the complement system, C4b-binding protein (C4BP) (5–7). For the alternative pathway of the complement system, the same function as C4BP is provided by factor H (FH, 150 kDa) (8, 9). FH binds C3b and thereby prevents formation of the alternative pathway proteolytic complexes C3(H_2_O)Bb and C3bBb, which produce C3b. FH also promotes the dissociation of C3(H_2_O)Bb and C3bBb, and acts as a cofactor for the complement factor I-mediated proteolytic cleavage and inactivation of C3b. Unique among the three complement pathways, the alternative pathway is constitutively active and has the potential for exponential amplification.

The M protein is one of the main Strep A recruiters of FH (8, 10, 11). This dimeric α-helical coiled-coil protein forms a dense fibrillar coat on the Strep A surface (12–14). Its surface location makes the M protein a target of immune surveillance, explaining its considerable antigenic sequence variation. The N-terminal 50 amino acids of the mature form of the M protein (i.e., after removal of the signal sequence) are hypervariable and define the M type. More than 250 distinct M protein types have been identified (15). Along with the M type-defining N-terminal hypervariable region, the remaining N-terminal third to half of M proteins is also sequence variable, while the C-terminal portion is conserved. The variable regions of M5, M6, and M18 proteins have been shown to directly bind FH (10, 11). However, due to sequence variability, no consensus FH-binding sequence motif is apparent in these M proteins. FH is composed of 20 complement control protein (CCP) domains, which are typically ∼60 amino acid β-sandwich folds containing two disulfide bonds. M5 and M6 proteins require FH domain 7 for binding (16, 17), which along with domains 6 and 8 interacts with human cell glycosaminoglycans (GAGs) (18, 19). FH domains 6-8 are also targeted by other pathogens, including *Neisseria meningitidis* and *Borrelia burgdorferi* (20, 21). Occupancy of these FH domains leaves domains 1-4 free to bind C3b and exert a negative regulatory role (22).

The M protein is also the major Strep A recruiter of C4BP, and as with FH, C4BP is bound by sequence variable regions of M proteins. The basis for M protein interaction with C4BP was elucidated through structural studies (6, 7). The variable regions of numerous M proteins present a set of identical or chemically conserved amino acids from their coiled coils that interact with a common set of C4BP amino acids. The primary sequence positions of these amino acids within the coiled coil heptad repeat define a C4BP-binding sequence pattern. Four such C4BP-binding patterns were identified for M proteins, encompassing 87 M proteins (6, 7).

We sought to understand whether M protein variable regions also have conserved sequence patterns for binding FH. To this end, we determined the structures of M5 and M6 protein fragments bound to FH domains 6 and 7, or FH(6–7), and carried out mutagenesis studies of interacting M protein amino acids. This enabled the construction of separate FH-binding sequence patterns for M5 and M6 proteins, and identification of a set of M proteins that have these sequence patterns. In addition to M proteins, FH is recruited to the Strep A surface by FbaA, which is encoded in the *mga* regulon of numerous strains of differing M types (23–26). FbaA binds FH domain 7 (23) and also fibronectin (24). The FH-binding region of FbaA was suggested to form a coiled coil (27), suggestive of M protein mimicry. Thus, we also determined the structure of the FH-binding portion of FbaA in complex with FH(6–7) and validated the structure through mutagenesis. FbaA formed a three-helix bundle preceded by an extensive loop instead of a coiled coil. Although FbaA is allelically sequence variable, its critical FH-contacting amino acids were absolutely conserved. Including both M proteins and FbaA, we identified FH-binding sequence patterns in about half of the approximately 250 known Strep A strains.

## Results

### Structure of M5-Factor H complex

We first confirmed direct interaction between intact M5 and M6 proteins with intact FH (Fig. S1), and then cocrystallized M5 and M6 protein fragments with FH(6–7).

The M5 protein fragment, composed of amino acids (aa) 94-175, yielded cocrystals that diffracted to 2.25 Å resolution limit (Fig. 1, Table S1). The structure, which was determined by molecular replacement (Fig. S2A), revealed two nearly identical M5/FH(6–7) complexes (rmsd 0.24 Å) in the asymmetric unit of the crystal (Fig. S3A). Each complex contained two FH(6–7) molecules bound to an M5 dimer, thus affording four views of the M5-FH interface. The M5 protein dimer formed a canonical coiled coil (28, 29), and FH(6–7) resembled conformations seen when bound to *Neisseria* fHbp and *B. burgdorferi* CspZ (average rmsd 1.48 Å and 1.02 Å, respectively) (20, 21) (Fig. S4), or the GAG analog sucrose octasulfate (SOS) (19) (average rmsd 0.75 Å). While the M5/FH(6–7) interface was in well-defined electron density, parts of FH(6–7) distal to the interface were in weak electron density. The buried surface area was extensive (average 896 Å^2^ for M5 protein and 895 Å^2^ for FH), with the bulk of contacts made to FH7, which was positioned closer to the C-terminal end of the M5 protein coiled coil. The shape fit between M5 protein and FH, as evaluated by shape complementarity was moderate (average of 0.64 across the four interfaces, with 1.0 a perfect fit) (30).

**Figure 1.**
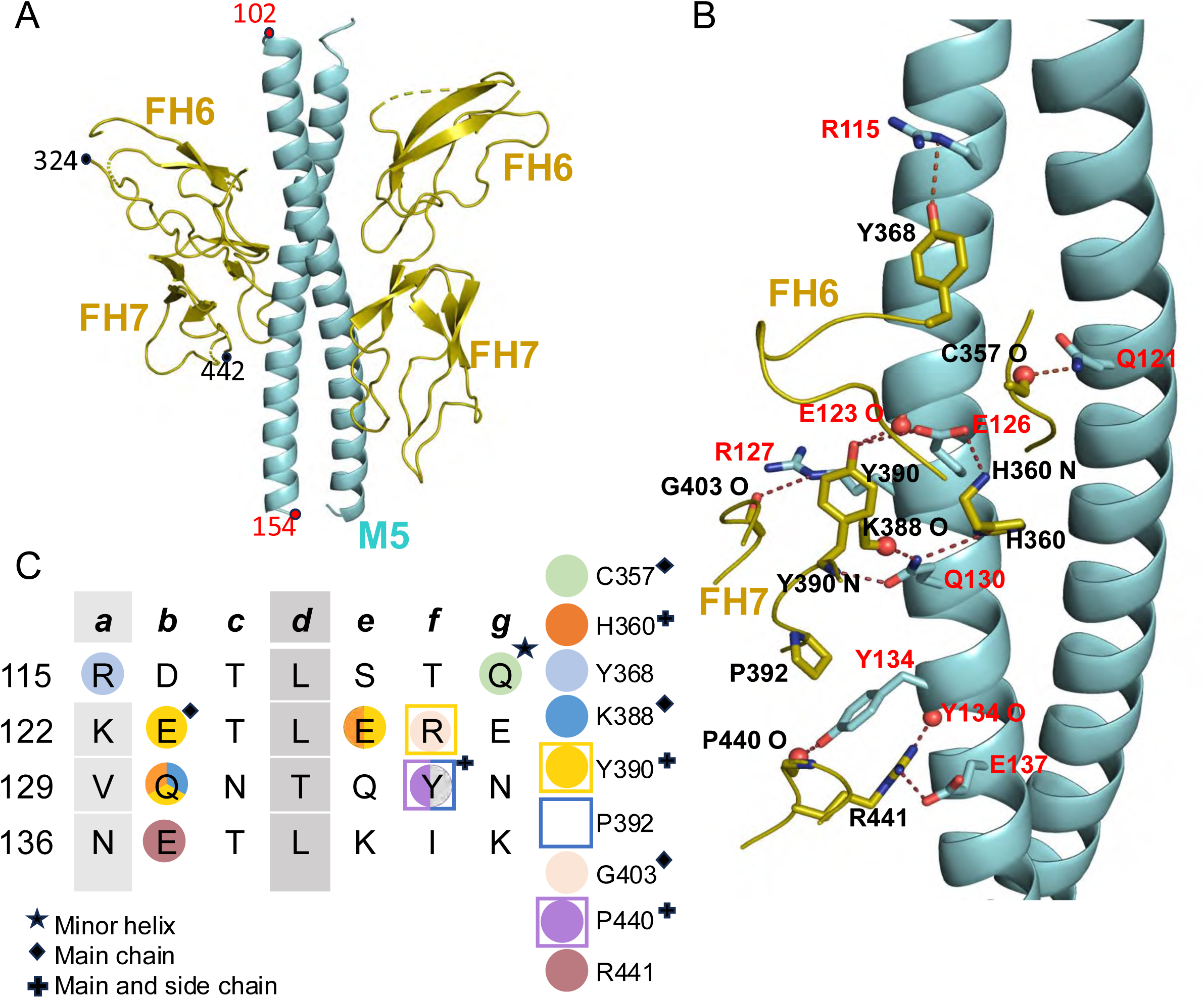
M5-FH interactions. **A.** Structure of M5 protein (cyan) in complex with FH(6–7) (olive), in cartoon representation. **B.** Interacting M5 protein and FH(6–7) amino acids. Red and blue spheres denote main chain carbonyls and amides, respectively. **C.** Heptad register of M5 protein (*a* and *d* positions in gray columns). Shapes indicate type of contact (circle for polar, square for hydrophobic), and coloring indicates which FH(6–7) amino acid (depicted in legend to the right) was contacted.

The interface between M5 protein and FH(6–7) was mostly polar and notable in consisting of a large number of contacts formed by main chain atoms of FH (Fig. 1). Each FH(6–7) molecule was contacted almost exclusively by a single α-helix of the M5 protein coiled coil. The FH-binding site in M5 protein spanned slightly more than four heptads (aa 115-137), with interactions concentrated in the central two heptads. The following interactions were observed in at least three of the four independent interfaces, unless otherwise noted. The second heptad supplied E126, which formed hydrogen bonds to the main chain amide of FH6 H360 and side chain of FH7 Y390; and R127, which formed a hydrogen bond to the carbonyl of FH7 G403, while its alkyl side chain was buried against the aromatic ring of FH7 Y390. In this second heptad, the main chain carbonyl of the M5 amino acid in the *b* heptad position (E123) formed a hydrogen bond to FH7 Y390. The third heptad supplied Q130, which in two of the interfaces formed hydrogen bonds to FH6 H360, and the main chain carbonyl and amide of FH7 K388 and Y390, respectively. The third heptad also supplied Y134, which formed a π-cation interaction with FH7 R441, a hydrogen bond with the main chain carbonyl of FH7 P440, and was positioned in a hydrophobic pocket formed by this proline and a second one, FH7 P392. The main chain carbonyl of Y134 also formed a hydrogen bond with FH7 R441. These interactions were preceded in the first heptad by one formed by R115, which is at the *a* heptad position and therefore destabilizing to the coiled coil. In two of the interfaces, M5 R115 formed a hydrogen bond with FH6 Y368. The first heptad also supplied the only contact from the minor α-helix, which was from Q121 and consisted of a hydrogen bond to the main chain carbonyl of FH6 C357. The fourth heptad supplied E137, which in two of the interfaces formed a salt bridge with FH7 R441.

The contribution of M5 amino acids to FH-binding was assessed by substitution mutagenesis. Purified intact wild-type or mutant M5 proteins were coated in ELISA wells and assayed for binding to soluble intact FH (Fig. 2A). Ala-substitution of three M5 amino acids had a profound effect in eliminating or nearly eliminating FH-binding. These were Q121 from the first heptad, E126 from the second, and Y134 from the third. Lys-substitution of E126 and the double E126A/Y134A substitution also eliminated FH-binding. By comparison, Ala-substitution of M5 R127 from the second heptad and Q130 from the third reduced interaction with FH moderately, to about one-third of wild-type M5 protein. Ala-substitution of M5 E137 from the fourth heptad decreased FH-binding by half, while Ala-substitution of M5 R115 from the first heptad increased FH-binding. The main chain of M5 E123 formed a contact with FH, but in the early stages of structure refinement, it appeared possible that its side chain would form a salt bridge with FH7 K388. Therefore, we constructed an M5 E123A mutant protein and found that its FH-binding was decreased by a quarter. Thus, it is possible that E123 forms a transient salt bridge with FH. We asked whether changes in FH-binding were due to non-specific alterations in the structure or stability of M5 mutant proteins. For this, circular dichroism spectra were collected as were temperature melting curves at 222 nm. Wild-type and mutant M5 proteins did not differ significantly in their CD spectra and stabilities (Fig. S5).

**Figure 2.**
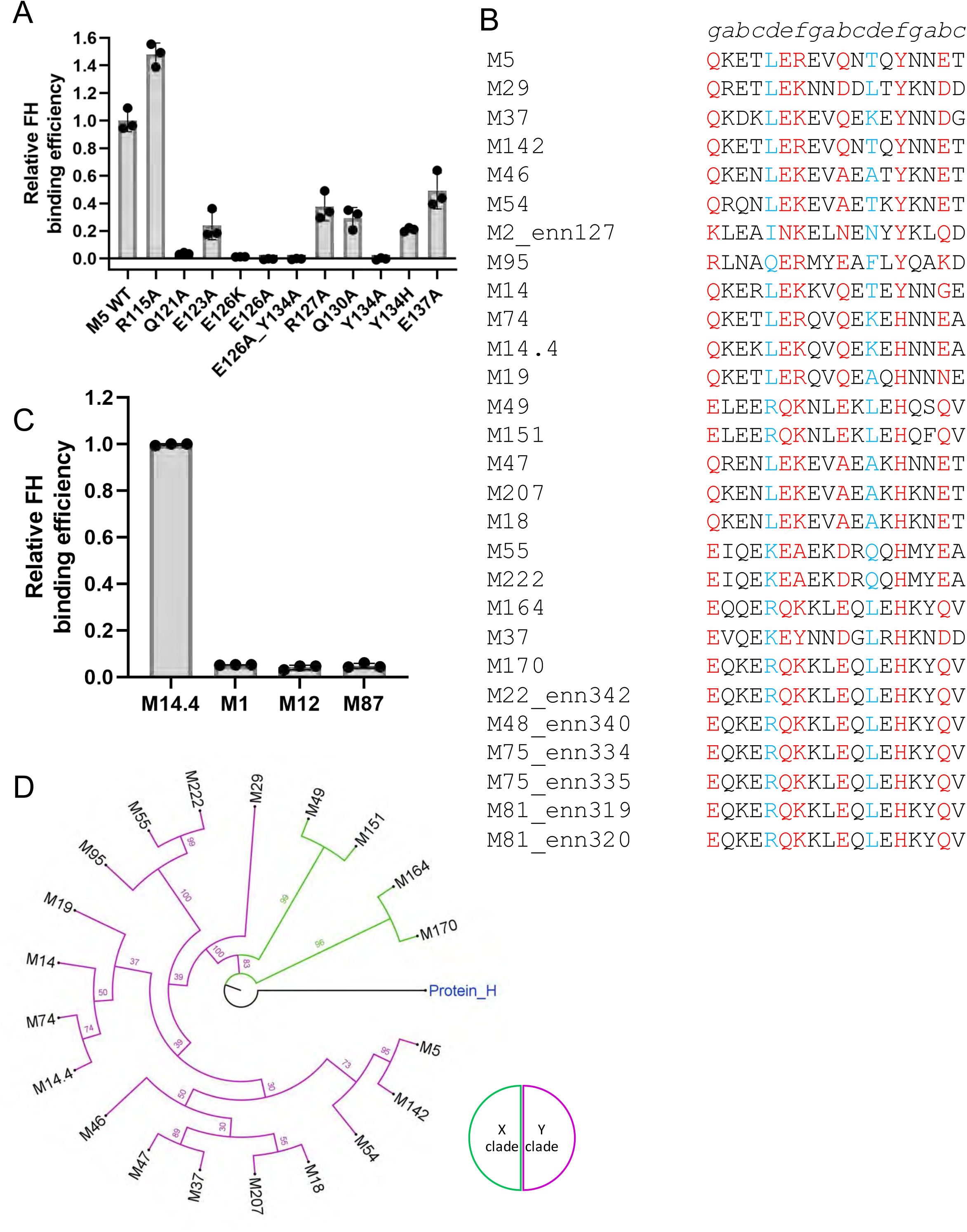
M5 protein FH-binding pattern. **A**. Binding of soluble intact FH protein to immobilized intact wild-type (WT) or mutant His_6_-M5 protein, as evaluated by ELISA. Bound FH was detected with an anti-FH monoclonal antibody. Values were normalized by FH binding to WT His_6_-M5 protein. Data from three biological replicates are presented with means and standard deviations. **B.** Sequences of M and Enn proteins with M5 FH-binding pattern. Heptad positions are indicated above the sequences. Red indicates an observed (M5 protein) or potential (other M and Enn proteins) FH-binding amino acid, and blue indicates a *d-*heptad position. M5 protein was placed at the top for reference. Other proteins are rank-ordered (Table S2). **C**. Binding of soluble intact FH to immobilized intact His_6_-M14.4, His_6_-M1, His_6_-M12, and His_6_-M87, evaluated as in panel A. Values were normalized by FH binding to His_6_-M14.4 protein. Data from three biological replicates are presented with means and standard deviations. **D.** Phylogenetic analysis of M proteins with M5 FH-binding pattern. Protein H was used as an outgroup.

### M5 FH-binding sequence pattern

We next asked if other M proteins had sequences that resembled the FH-binding sequence in M5 protein. A collection of 186 M protein sequences was scored. Those that had identical or chemically similar amino acids corresponding to the essential M5 amino acids Q121 (*g* heptad position), E126 (*b*), and Y134 (*f*) received high scores (Fig. 2B, Table S2). Sequences with identical amino acids received a higher score than ones with chemically similar ones, and sequences lacking identical or chemically similar amino acids at any of these positions were eliminated. The proper heptad position of these amino acids was also scored. Due to the often irregular heptads of M proteins, for example M5 R115 and K122 at *a* positions, only half of the *a* and *d* positions in three consecutive heptads needed to have coiled coil-stabilizing amino acids. A restriction was also imposed that these three heptads have no more than two coiled coil-destabilizing amino acids at *a* or *d* positions. Since Ala-substitution of M5 E123, R127, Q130, and E137 reduced but did not eliminate FH-binding, the occurrence of identical or chemically similar amino acids received positive but lower scores than the essential M5 amino acids.

The scoring yielded M18 protein, which was confirmed to bind FH (Fig. S1) (11). Since most M proteins identified through this scoring have not been experimentally assessed for FH binding, we examined several by ELISA (Figs. 2C and S1). We found that M14.4 protein, which scored higher on the list than M18 protein (Table S2), bound FH while M12 and M87 proteins, which scored lower than M18, did not. M1 protein, which does not bind FH, was used as a negative control in this experiment. Since M87 protein was the highest scoring non-binder, its score was used as a cut-off, which resulted in a total of 20 M proteins, including M5 protein, as having an M5 pattern for FH binding (Table S2). This includes two distinct isolates of M14 protein (M14 and M14.4, 76% identical). Interestingly, M37 protein had two overlapping FH-binding sequence patterns (Table S2). M proteins are assigned to two clades, X and Y (31), and 16 of these 20 M proteins belong to clade Y while 4 belong to clade X (Fig. 2D).

Many Strep A strains express M-like coiled-coil proteins in addition to M proteins. These are Enn proteins, which are sequence variable but less so than M proteins, and M-related protein (Mrp), which are also sequence variable but even less so than Enn proteins (32, 33). Collections of 144 Enn proteins and 221 Mrps were searched for sequences that matched the FH-binding sequence in M5 protein, with the score for M87 protein serving as a cut-off. No Mrp was identified but one Enn protein (Enn127 from an M2 strain) had a higher score than M18 protein, suggesting that it is capable of binding FH. There were also six other Enn proteins belonging to four M types (M22, M48, M75, and M81) that scored above the M87 protein cut-off and may have the capability of binding FH.

### Structure of M6-Factor H complex

An M6 protein fragment composed of amino acids 74-157 yielded cocrystals with FH(6–7) that diffracted to 1.90 Å resolution limit (Fig. 3, Table S1). The structure, which was determined by molecular replacement (Fig. S2B), contained two nearly identical complexes (rmsd 1.49 Ả) in the asymmetric unit of crystal (Fig. S3B). Each complex contained an M6 dimer but only one FH(6–7) molecule. The presence of one rather than two FH molecules appears to be due to the second, dimer-related FH-binding site in M6 protein being blocked by a crystal contact to another M6 protein dimer (Fig. S6). The M6 protein dimer formed a canonical coiled coil, and FH(6–7) was structurally similar to SOS-, fHbp-, CspZ-bound forms (average rmsd of 0.80 Å, 1.12 Å, 0.78 Å, respectively). The interface of the complex was in well-defined electron density, while some parts of FH(6–7) distal to the interface had weak electron density. The buried surface area was extensive for the M6/FH(6–7) complex (average 751 Å^2^ for M6 and 653 Å^2^ for FH), but unlike the M5/FH(6–7) complex, the two M6 α-helices contributed almost equally to the binding of a single FH(6–7) molecule. The majority of contacts were made to FH6 by the two M6 α*-*helices (called M6a and M6b). A few positions in FH7 were also contacted by M6b. The shape fit between M6 and FH, as evaluated by shape complementarity, was high (average of 0.708).

**Figure 3.**
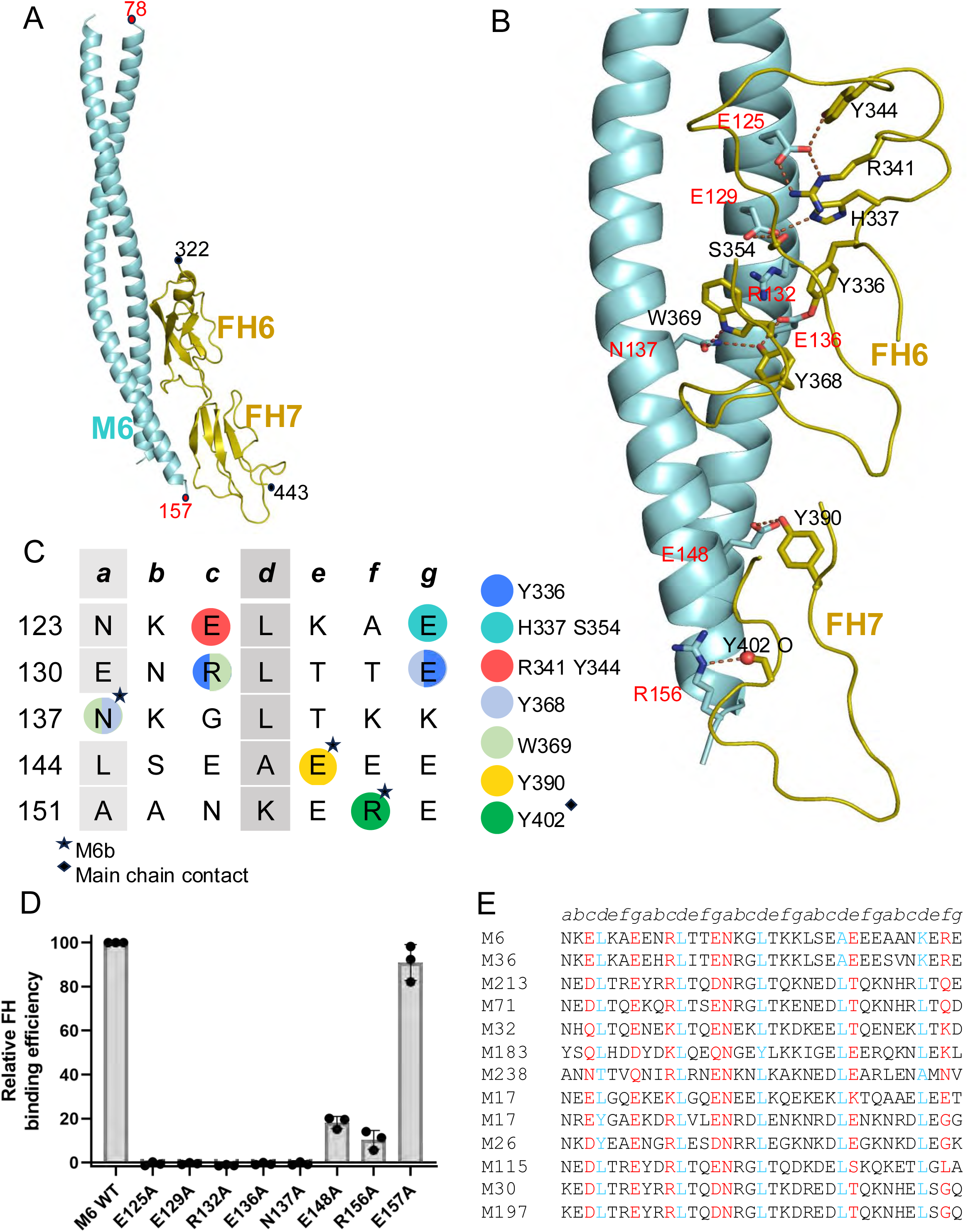
M6-FH interactions. **A**. Structure of M6 protein (cyan) in complex with FH(6–7) (olive), in cartoon representation. **B.** Interacting M6 protein and FH(6–7) amino acids. Red and blue spheres denote main chain carbonyls and amides, respectively. **C.** Heptad register of M6 protein (*a* and *d* positions in gray columns). Circles indicate polar contacts, and coloring indicates which FH(6–7) amino acid (depicted in legend to the right) was contacted. **D.** Binding of soluble, intact FH protein to immobilized intact wild-type and mutant His_6_-M6 protein, as evaluated by ELISA. Bound FH was detected with an anti-FH monoclonal antibody. Values were normalized by FH binding to wild-type His_6_-M6 protein. Data from three biological replicates are presented with means and standard deviations. **E.** Sequences of M proteins with M6 FH-binding pattern. Heptad positions are indicated above the sequences. Red indicates an observed (M6 protein) or potential (other M protein) FH-binding amino acid, and blue indicates *a d-*heptad position. The list is rank-ordered (Table S2).

The FH-binding site in M6 spanned five heptads (aa 125-156) (Fig. 3), and involved primarily aromatic amino acids of FH(6–7) rather than the main chain as in the M5 protein complex. In the first heptad, M6a E125 formed a salt bridge with FH6 R341 and a hydrogen bond with FH6 Y344, and M6a E129 formed hydrogen bonds with FH6 H337 (or a salt bridge, depending on the protonation state of the His) and FH6 S354. In the second heptad, M6a R132 formed π-cation interactions with FH6 Y336 and W369, and M6a E136 formed hydrogen bonds with FH6 Y336 and Y368. The next three heptads provided three contacting amino acids from the M6b α-helix: M6b N137 formed a hydrogen bond with FH6 W369 but also had the potential to form a hydrogen bond with FH6 Y368, accompanied by a flip of the Asn side chain; M6b E148 formed a hydrogen bond to Y390; and M6b R156 formed a hydrogen bond with the main chain carbonyl of FH7 Y402.

Ala-substitution of all FH-contacting amino acids in M6 protein, except for E148 and R156, resulted in complete loss of FH-binding (i.e., E125A, E129A, R132A, E136A, N137A) (Fig. 3D). Binding was measured by ELISA as described for M5 protein. Approximately 10-20% of FH-binding was maintained in M6 E148A and R156A. Due to the large number of Ala-substitutions that resulted in complete loss of binding, we constructed an Ala-substitution at M6 E157, an amino acid that is not observed to contact FH(6–7). M6 E157A maintained FH-binding at wild-type level, providing validation for this assay. The five mutant M6 proteins that were entirely deficient in FH-binding had CD spectra resembling that of wild type M6 protein (Fig. S7). The stability of these mutants was either not significantly different from (M6 E136A and N137A) or exceeded (E125A and E129A) that of wild-type M6 protein, while loss of FH-binding in M6 R132A was at least partially attributable to a statistically significant loss in stability (Fig. S7).

### M6 FH-binding sequence pattern

As for M5 protein, we scored M protein sequences for a pattern of FH-contacting amino acids resembling that of M6 protein (Fig. 3E, Table S3). A stretch of five heptads was scored for at least four stabilizing amino acids in *a* or *d* positions, and no more than two destabilizing amino acids at these positions. This scoring excluded the *a* position of the third heptad, since this *a* position corresponds to the FH-contacting amino acid M6 N137. For the five functionally essential M6 amino acids (E125, E129, R132, E136, and N137), identical or chemically conserved amino acids were required. At the two positions equivalent to E148 and R156, at which Ala-substitution diminished but did not eliminate FH binding, the occurrence of identical or chemically similar amino acids received a positive score, but sequences were not eliminated if other amino acids occurred at these positions. A rank-ordered list of M proteins was produced, which included the FH non-binder M87 protein (Table S3). Thus, the score of M87 protein was used as an initial threshold. No Mrps scored higher than M87 protein, but some Enn proteins did. One of the highest scoring Enn proteins, Enn300, along with another high scoring Enn protein, Enn262, were assayed for FH-binding by ELISA and found not to bind (Fig. S8). Thus, the score for Enn300, which was higher than that for M87 protein, was set as the threshold, which resulted in the identification of a total of 12 M proteins, including M6 protein, as having an M6 pattern for FH binding (Table S3). Two tandem M6 FH-binding patterns occurred in M17 protein. Eleven of the 12 M proteins belong to clade Y and only M183 belongs to clade X (Fig. S8).

### Structure of FbaA/Factor H complex

The FH-binding region of FbaA (aa 57-130) was cocrystallized with FH(6–7) (27). The crystal structure of the complex was determined by molecular replacement to 1.82 Å resolution limit (Figs. 4 and S2C, and Table S1). Two nearly identical complexes (rmsd 0.36 Å) were present in the asymmetric unit of the crystal (Fig. S3C). The FbaA fragment was monomeric and formed a loop (aa 57-72) followed by a three-helix bundle (aa 73-124) instead of a coiled coil. AlphaFold3 predicts a three-helix bundle as well (34). The FbaA fragment was determined by SEC-MALS to be monomeric, as observed in the crystal structure, rather than dimeric, as would occur in an M protein-like coiled coil (Fig. S9). FH(6–7) was unchanged from its structures when bound to M5 protein, M6 protein, SOS, *Neisseria* fHbp, or *Borrelia* CspZ (1.07, 1.68, 1.06, 2.15, and 1.55 Å rmsd, respectively). The FbaA-FH(6–7) interface was in well-defined electron density, and the buried surface area was extensive (average 776 Å^2^ for FbaA and 692 Å for FH(6–7)). The shape fit between FbaA and FH, as evaluated by shape complementarity, was high (average of 0.731).

**Figure 4.**
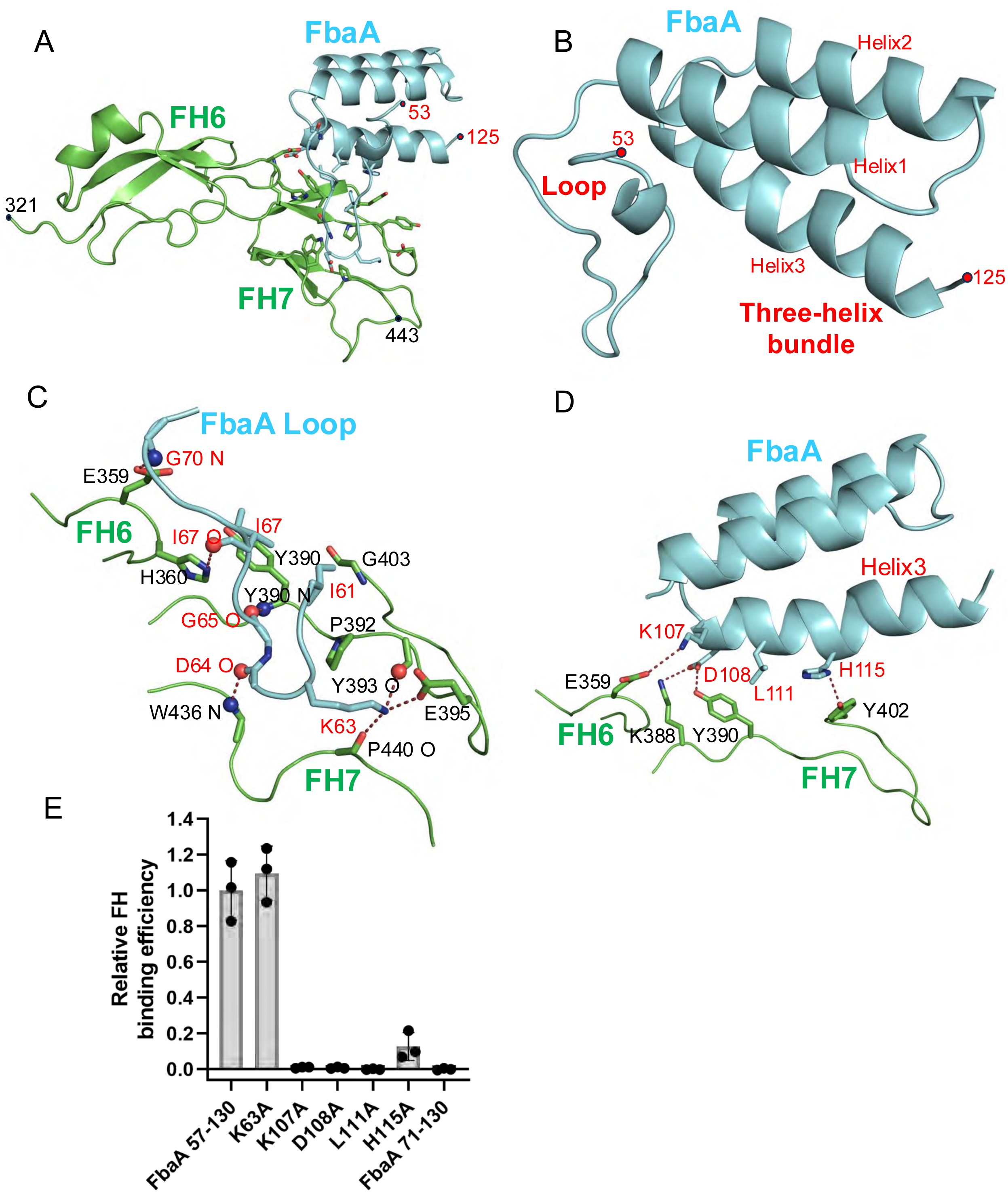
FbaA-FH interactions. **A.** Structure of FbaA (57-130, cyan) in complex with FH(6–7) (green), in cartoon representation. **B.** Structure of the loop and three-helix bundle of FbaΑ (57–130). **C.** Interaction between FbaA(57–130) loop and FH(6–7). Red and blue spheres denote main chain carbonyls and amides, respectively. **D.** Interaction between the FbaA (57–130) three-helix bundle and FH(6–7), with spheres as in panel C. E. Binding of soluble intact FH to immobilized wild-type and mutant His_6_-FbaA (57–130), as evaluated by ELISA. Bound FH was detected with an anti-FH monoclonal antibody. Values were normalized by FH binding to wild-type FbaA. Data from three biological replicates are presented with means and standard deviations.

The N-terminal loop of FbaA along with the three-helix bundle mainly contacted FH7, with most of these contacts being polar and many involving main chain atoms of FbaA and FH (Fig. 4). The FbaA loop supplied the side chain of I61 (Fig. 4C), which was buried in a hydrophobic pocket formed by FH7 P392, Y390, and the Cα of G403; K63, which formed hydrogen bonds to the main chain carbonyls of FH7 Y393 and P440, and in one interface, a salt bridge with FH7 E395; and I67 which formed a hydrophobic contact with FH7 Y390. Main chain atoms from the FbaA loop formed several contacts: the carbonyls of FbaA D64 and G65 formed hydrogen bonds to the amides of FH7 W436 and FH6 Y390, respectively; the carbonyl of I67 formed a hydrogen bond to FH6 H360; and the amide of FbaA G70 formed a hydrogen bond with FH6 E359. In the three-helix bundle, the third helix contributed the bulk of contacts (Fig. 4D). FbaA K107 formed a salt bridge with FH6 E359. FbaA D108 formed a hydrogen bond and salt bridge with FH7 Y390 and K388, respectively. FbaA L111was buried against FH7 Y390, and FbaA H115 formed a hydrogen bond with FH7 Y402.

We determined the contribution to FH-binding of several FbaA side chains. Ala-substitution of K107 and D108 in FbaA (57–130) eliminated FH-binding, and that of H115 reduced FH-binding by ∼10-fold compared with wild-type FbaA (57–130) (Fig. 4E). Deletion of the FbaA loop (Δ57-70) also eliminated FH-binding. In contrast, Ala-substitution of K63 left FH-binding unchanged. FbaA K107A was slightly altered in structure, as assessed by CD, whereas the other mutations, including the loop deletion, had similar circular dichroism spectra (Fig. S9). The stabilities of mutant FbaA (57–130), even with the slight structural alteration in K107A, were not significantly different from that of wild-type.

The sequences of a collection of global Strep A isolates composed of 148 M types was searched for FbaA sequences (32), with 95 strains containing such sequences (Fig. 5). The M types were distributed fairly evenly between X and Y clades (46 and 49, respectively). The entirety of the FbaA sequence (∼400 aa) had an average identity of 80% (range 73-90%), while the FH-binding loop and three-helix bundle region had an average identity of 73% (range 56-100%). Notably, six of the seven FbaA amino acids identified to contact FH through their side chains were absolutely conserved (Fig. S10). The only one not absolutely conserved was K63, which was relatively unimportant for FH-binding (Fig. 4E). Some of the M types that contained FbaA had in addition an M protein with a M5 FH-binding pattern (M14.4, M74, M95) or M6 FH-binding pattern (M71 and M115), an Enn protein with an M5 FH-binding pattern (M22, M75, and M81), or both an M and Enn protein with an M5 FH-binding pattern (M2).

**Figure 5.**
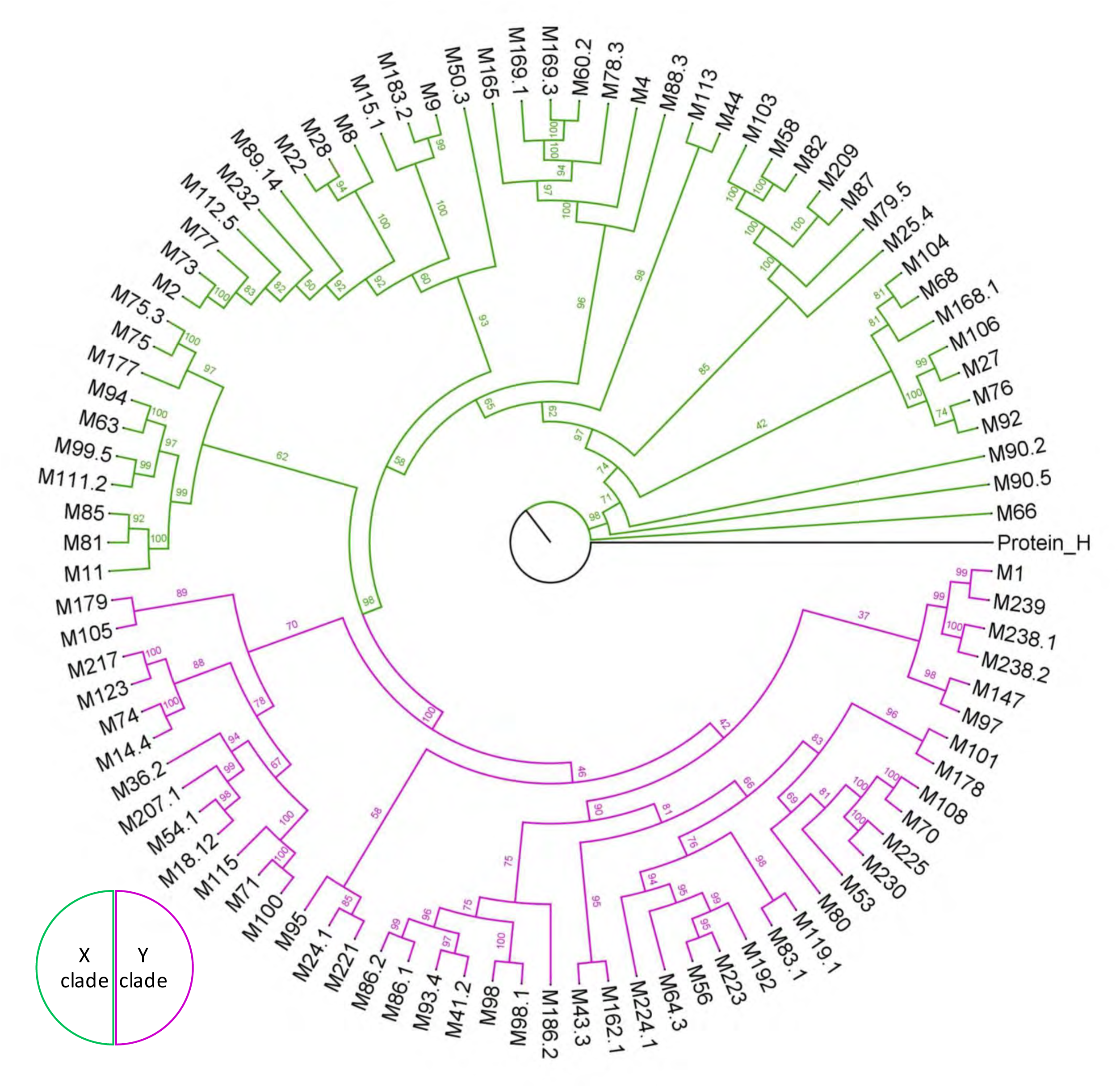
Phylogenetics of FbaA. Phylogenetic analysis of the M types of FbaA-expressing Strep A strains. Protein H was used as an outgroup.

## Discussion

We determined the mechanisms by which M5 protein, M6 protein, and FbaA bind human FH. While all three proteins contacted FH domains 6 and 7, their modes of interaction differed considerably. M5 protein and FbaA were alike in primarily contacting FH7 whereas M6 protein primarily contacted FH6. In the case of M5 protein and FbaA, main chain contacts were noteworthy. Numerous main chain atoms of FH7 were contacted by either side chain atoms of M5 protein or main chain atoms of FbaA, reflecting shape fits in these two proteins to the FH7 domain backbone. In contrast, M6 protein primarily contacted aromatic amino acids of FH6. A polymorphism associated with age-related macular degeneration in FH7, Y402H, is reported to decrease interaction with certain Strep A strains (35, 36). While M5 protein did not contact Y402 and M6 protein contacted only the main chain of Y402, FbaA H115 formed a hydrogen bond to the side chain of Y402, suggesting that FbaA is likely to be responsible for FH-binding in these strains. In addition, FH7 R369A/K370A and R386A/K387A are reported to decrease binding to M6 protein (37). These FH amino acids are proximal but not in direct contact with M6 protein, and thus the decreased binding is likely to be an indirect effect.

Only one interacting FH amino acid, Y390, was common to M5 protein, M6 protein, and FbaA, in all cases being contacted by a glutamate. FH Y390 is also contacted by GAGs (19), *Neisseria* fHbp (20), and *B. burgdorferi* CspZ (21) (Figs. S11-S13). *Neisseria* fHbp mimics GAGs in its interaction with FH(6–7), with numerous contacted FH amino acids in common. M6 protein mimics neither GAGs nor fHbp, but half or more of the FH amino acids it contacts are also contacted by GAGs or fHbp. CspZ, like M5 protein and FbaA, interacts primarily with FH domain 7, and half or more of the FH amino acids contacted by M5 protein or FbaA are also contacted by CspZ. These observations indicate that FH domains 6 and 7 are interaction hot spots, and that they have a remarkable ability to accommodate different binding modes.

The M5 and M6 FH-binding sequence patterns are distinct and define two families of FH-binding M proteins. The M5 FH-binding pattern was identified in a total of 20 M types after application of experimentally determined scoring cut-offs, and the M6 sequence pattern in a total of 12 M types. The M6 sequence pattern was especially sensitive to loss of function through mutation, as Ala-substitutions at five of the seven FH-contacting amino acids eliminated FH binding. Two M proteins with the M5 FH-binding sequence pattern, M14 and M18, were demonstrated to bind FH directly. In addition, several M proteins with the M5 FH-binding sequence patterns belong to strains demonstrated to bind FH (i.e., M29, M55, M74, and M95) (38, 39). In the case of the M6 FH-binding sequence pattern, only M6 protein has been experimentally verified to bind FH. However, it is notable that two of the M proteins identified as having an M6 FH-binding sequence patterns, M30 and M71, are from strains demonstrated to bind FH (38, 39). The M5 FH-binding pattern was also identified in one Enn protein with a high score and five others with scores just above the cut-off. Whether these Enn proteins bind FH requires experimental testing.

The 32 M proteins and up to 6 Enn proteins identified to have a FH-binding pattern represent 37 M types, which is much fewer than the 87 M and 113 Enn proteins identified to have C4BP-binding patterns representing 111 M types (6, 7). FH- and C4BP-binding generally assort to different M protein clades, Y and X, respectively. Only two M proteins have both FH- and C4BP-binding patterns, the X clade M49 protein and Y clade M95 protein. Unlike C4BP, FH is bound by Strep A proteins other than M and Enn proteins, most prominently FbaA. We found that the FH-binding segment of FbaA consists of a loop and three-helix bundle rather than a coiled coil. The FbaA three-helix bundle occurs in the midst of intrinsically disordered regions (34), some of which are likely form tandem β-zippers that interact with fibronectin (40). Notably, while FbaA displays allelic variation, the amino acids important to FH-binding were identical across alleles. Ninety-five strains of differing M types encoded FbaA, with nearly equal apportionment between X and Y clades. With these 95 strains and those that have either M or Enn proteins with FH-binding sequence patterns, we tabulated a total of 123 Strep A strains of differing M types (including subtypes) that have the capacity to bind FH, very similar to the representation of C4BP-binding capacity in 111 Strep A strains (7). This conclusion is consistent with a survey showing that 24 out of 38 strains of differing M types bind FH (39). C4BP- and FH-binding occur together in numerous X clade strains, as C4BP-binding is concentrated in the X clade and FbaA is distributed across both clades. While extensive evidence exists for the importance to virulence of C4BP recruitment (41–43), the evidence for FH recruitment is less certain. For example, FH recruitment is found to have little or no effect on bacterial resistance to killing in human blood (11, 44, 45), and evidence as to whether C3b deposition is affected by FH recruitment is contradictory (8, 10, 45). The prevalence of FH-binding in Strep A strains, mainly due to FbaA, strongly argues for FH recruitment having a role in virulence. The mechanistic insights into FH interaction provided by our structural and functional experiments are applicable to precise investigation of the significance of FH recruitment to the Strep A surface.

## Experimental Methods

### Bacterial strains and DNA manipulation

The coding sequences of mature M5 protein (aa 43–377, strain Manfredo, gift from P. Nordenfelt, Lund University, Sweden) and M6 protein (aa 43-452, strain JRS4, gift from T. Barnett, Telethon Kids Institute, Perth, Australia) were cloned into the vector pET28b (Novagen), with a C-terminal His_6_-tag encoded. Coding sequences for truncated forms of M proteins and FbaA (aa 57-130, M1 strain 5448) were cloned into a modified version of pET28a vector (Novagen), which encoded an N-terminal His_6_-tag followed by a PreScission™ protease cleavage site. The coding sequence for FH domains 6-7 were chemically synthesized with codons optimized for *E. coli* expression (IDT), and cloned into a modified pET28a vector. Amino acid substitutions and deletions were introduced into pET28b vectors with the QuickChange II Site-Directed mutagenesis kit (Stratagene), according to the manufacturer’s directions.

### Protein Expression and Purification

M proteins, Enn proteins and FbaA constructs were expressed in soluble form in *Escherichia coli* BL21 (Gold) and purified as previously described (4, 6). FH domains 6-7 (FH(6–7)) were expressed as inclusion bodies in *E. coli* BL21 (Gold) and refolded and purified as previously described (19).

### Protein quantification

Concentrations of purified M proteins were determined using Bradford Assay (Bio-Rad) according to manufacturer’s instructions. The concentrations of FH(6–7) and FbaA (57–130) concentration was measured using their A_280_ and calculated molar extinction coefficient of 36370 M^-1^ cm^-1^ or 9970 M^-1^ cm^-1^, respectively.

### Molecular mass determination

His-tagged FbaA (57–130) (2.0 mg/ml) in 20 mM Tris-Cl, 150 mM NaCl, pH 8.0 was centrifuged (10 min, 20,000 × *g*, 20 °C) to remove aggregates. The sample (100 μl) was then immediately applied to a Zenix SEC-300 column, which had been equilibrated in the same buffer, at RT. Samples eluting from the column were monitored with a light scattering detector (DAWN HELEOS II, Wyatt Technology, Santa Barbara, CA) and a differential refractometer (Optilab T-rEX; Wyatt Technology). Data processing and molecular mass calculation were performed with ASTRA software (Wyatt Technology).

### Crystallization and data collection

M5 protein (aa 94-175) and M6 protein (aa 74-157) constructs and FbaA (aa 57-130) were mixed with FH(6–7) at a 1:1 molar ratio, and dialyzed overnight at 4 °C in 50 mM Tris, 150 mM NaCl, pH 8. The samples were then concentrated by ultrafiltration using a 3,000 MWCO membrane to ∼10 mg/ml (Millipore; 4,500 x *g*, 30 min, 4 °C). Crystallization was performed by the hanging drop vapor-diffusion method. M5-FH(6–7), M6-FH(6–7), and FbaA-FH(6–7) were co-crystallized at 18 °C by mixing 1 μl of the complex with 1 μl of the reservoir solution. In the case of M5-FH(6–7), the reservoir solution was 20% PEG 6000, 15% ethylene glycol, and 0.2 M NaCl. For M6-FH(6–7), it was 30% PEG 1000, 0.1 M MMT (DL-malic acid:MES:Tris base), pH 7.5 and for FbaA-FH(6–7), 20% PEG 6000, 8% ethylene glycol, 0.2 M NaCl, and 0.1 M MES, pH 6.0. Crystals were transferred to their respective reservoir solutions supplemented with 30% ethylene glycol for cryopreservation, mounted in fiber loops, and flash-cooled in liquid N_2_.

Diffraction data for M5-FH(6–7) and M6-FH(6–7) were collected at the Advanced Photon Source (APS, beam line 22-ID) and Stanford Synchrotron Radiation Light source (SSRL, beamline 9-2), respectively. Diffraction data for FbaA-FH(6–7) were collected at the UCSD X-ray Crystallography facility. Diffraction data for M5-FH(6–7) were indexed, integrated, and scaled with HKL 2000 (46); data were truncated in the last resolution shell in which CC* was > 0.3 (47). For M6-FH(6–7), data were indexed, integrated (DIALS), and scaled (Aimless) with CCP4i2 (48), and truncated in the last resolution in which CC_1/2_ was > 0.5. For FbaA-FH(6–7), data were indexed and integrated using the Bruker SAINT Software program and scaled using SADABS software, and truncated as for M6-FH(6–7). Analysis of diffraction intensities with Phenix/Xtriage (49) indicated that the M5-FH(6–7) crystals were merohedrally twinned.

### Structure determination and refinement

Phases for the M5-FH(6–7) complex were determined by molecular replacement using FH domains 6 and 7 from the structure of FH domains 6, 7, and 8 (PDB: 2UWN) as the search model with Phaser (50). The asymmetric unit was identified to contain four FH(6–7) molecules. Continuous electron density corresponding to the M5 protein construct was evident in initial electron density maps. A second round of molecular replacement was carried out with the four FH(6–7) molecules fixed, and a dimeric, α-helical coiled coil model of M5 protein (aa 94-175) generated with ColabFold (51) as the search model. The model was built manually into well-defined electron density using Coot and modified as guided by the inspection of σA-weighted 2Fo-Fc and Fo-Fc omit maps. Refinement was performed using Refine from the Phenix suite (52) with default settings. Side chains with no electron density beyond Cβ were truncated at Cβ. Twinning was taken into account in phenix refinement using the twin law -h, -k, l.

Phases for the M6-FH(6–7) complex were determined by molecular replacement, as described for the M5-FH(6–7) complex. The asymmetric unit was identified to contain two FH(6–7) molecules. A second round of molecular replacement was carried out with the two FH(6–7) molecules fixed, and a dimeric, α-helical coiled coil model of M6 protein (aa 74-157) generated with ColabFold (51) as the search model. The model was built manually into well-defined electron density using Coot and modified as guided by the inspection of σA-weighted 2Fo-Fc and Fo-Fc omit maps. Refinement was performed using Refine from the Phenix suite with default settings. At late stages, translation-libration-screw (TLS) parameters were introduced (chain A; 5 groups, chain B; 6 groups, chain C; 3 groups, chain D; 2 groups, chain E; 3 groups and chain F; 4 groups). Side chains with no electron density beyond Cβ were truncated at Cβ.

Phases for the FbaA-FH(6–7) complex were determined as described for the M5-FH(6–7) complex. Two FH(6–7) molecules occupied the asymmetric unit of the crystal. A second round of molecular replacement was carried out with the two FH(6–7) molecules fixed, and an FbaA protein (74–157) model generated with ColabFold (51) as the search model. The placement of FbaA was inspected and modified manually in Coot as guided by the σA-weighted 2Fo-Fc and Fo-Fc omit maps. Refinement was performed using Refine from the Phenix suite with default settings. At late stages, TLS parameters were introduced (chain A and chain B constituted 2 TLS groups, whereas chain C and Chain D constituted 3 groups). Side chains with no electron density beyond Cβ were truncated at Cβ.

The interaction interfaces between protein complexes were analyzed with PISA (53) and visualized with Pymol. Pymol (Schrödinger, LLC) was used for generating molecular figures.

## ELISA

### Detection of Factor H binding

Purified mature M and Enn proteins at 20 μg/ml were coated on the wells of 96-well microtiter plates (Corning) in phosphate buffered saline (PBS, pH 7.2) overnight at 4 °C. All subsequent steps were performed at RT. Wells were washed three times in TBST (150 mM NaCl, 50 mM Tris, pH 8.0, and 0.1% Tween-20) and blotted dry between each step. Wells were blocked with 0.1% BSA in PBS for 1 h, and then incubated with 100 μl factor H (10 μg/ml, Complement Technology) for 1 h. After washing three times with TBST, 100 μl of mouse monoclonal anti-human FH antibodies (Biolegend) at 1:1000 dilution in 0.1% BSA/PBS were added to wells for 1 h. To detect bound FH, wells were washed with TBST three times, and 100 μl horseradish peroxidase (HRP)-conjugated goat anti-mouse IgG (Biolegend) at 1:1500 dilution in 0.1% BSA/PBS was added to the wells and incubated for 1 h. One hundred μl TMB substrate (BD Biosciences) was then added to the wells and incubated for 10 min (protected from light), followed by addition of 50 μl 2 N sulfuric acid to stop the reaction. The absorbance at 450 nm was measured. Uncoated wells treated in the same way were used as a background and subtracted from the binding measurements.

### CD spectroscopy

CD spectra were measured on an J1500 Circular Dichroism Spectrometer using a quartz cell with a 1 mm path length. Protein samples were ∼0.2 mg/ml in 5 mM Tris-HCl, pH 8.0. Wavelength spectra were recorded in a range of 190–260 nm at 25 °C at 0.1 nm intervals with a 1 sec averaging time per data point. Thermal melting was carried out between 20 and 60 °C in 1 °C increments, with the CD signal being monitored at 222 nm.

### Sequence Scoring

Files containing 186 M protein, 143 Enn protein sequences, and 221 Mrp sequences (from NCBI GenBank) (32) were searched for FH-binding patterns. The sequences were in fasta format, and a python script was written to carry out the search.

For sequence patterns corresponding to M5 protein, a stretch of 19 aa were searched. Only sequences that had coiled-coil stabilizing amino acids (i.e., V, L, I, M, F, or Y) at three or more *a* or *d* positions in three consecutive heptads, and no more than two coiled-coil destabilizing amino acids at the positions (H, K, D, S, R, or G at *a*, and the same at *d* in addition to T and N) were considered. At the functionally essential positions corresponding to M5 Q121, E126, and Y134, the occurrence of an identical or chemically conserved amino acid was required. For the position equivalent to Q121, identical and chemically conserved amino acids were given the same score of +1, and chemically conserved were those capable of hydrogen-bonding and being accommodated at the interface (as evaluated by modeling): Q, N, Y, R, K, D, E or W. For the position equivalent to E126, the occurrence of D or E resulted in a score of +1, whereas N or Q resulted in a score of +0.7. For the position equivalent to Y134, the occurrence of Y resulted in a score of +1, whereas H, F, or W resulted in a score of +0.2. Other positions were scored positively for the occurrence of specific amino acids, but sequences were not eliminated if no such amino acids were present at these positions. The scoring of Ala at these positions correlated with the relative loss of FH-binding due to Ala-substitutions. These positions were as follows. At the position equivalent to M5 E123, the occurrence of D or E resulted in a score of +0.5, N or Q a score of +0.3; and A a score of +0.25. At the position equivalent to M5 R127, the occurrence of R or K resulted in a score of +0.7, and A a score of 0.4. At the position equivalent to M5 Q130, the occurrence of D or E resulted in a score of +0.7, N or Q a score of +0.5, and A a score of +0.3. At a position equivalent to M5 E137, the occurrence of D or Ε resulted in a score of +0.7, N or Q a score of +0.6, and A a score of +0.5.

For sequence patterns corresponding to M6 protein, a stretch of 35 aa were searched. Only sequences that had coiled-coil stabilizing amino acids (i.e., V, L, I, M, F, or Y) at four or more *a* or *d* positions in five consecutive heptads, excluding the equivalent of M6 N137 at the *a* position, and no more than two coiled-coil destabilizing amino acids at the positions (H, K, D, S, R, or G at *a*, and the same at *d* in addition to T and N) were considered. At the functionally essential positions corresponding to M6 E125, E129, R132, E136, and N137, the occurrence of an identical or chemically conserved amino acid was required. For the positions equivalent to E125, E129, and E136, the D or E was scored +1, and N or Q +0.7. For the position equivalent to R132, the occurrence of R or K was scored +1. For the position equivalent to N137, the occurrence of N was scored +1 and Q +0.7, and D or E +0.4. This scoring was based on the fact that N137 occupies an *a* heptad repeat position. Positions equivalent to the M6 amino acids at which Ala-substitution diminished but did not eliminate FH-binding, E148 and R156, were scored positively for the occurrence of specific amino acids, but sequences were not eliminated if no such amino acids were present at these positions. The scoring of Ala at these positions correlated with the relative loss of FH-binding due to Ala-substitutions. For the position equivalent to E148, the occurrence of D or E was scored +0.7, N or Q +0.5, and A +0.2. For the position equivalent to R156, the occurrence of R, K, D, E, N, or Q was scored +0.7 and A +0.1.

### Phylogenetic analysis of FH binding *S. pyogenes* strains

Multiple protein sequence alignments of M proteins were carried out using MUSCLE in MEGA11 with default parameters (54, 55). IQ-TREE v1.6.12 was used for the generation of maximum likelihood (ML) tree of 186 M proteins under the substitution model (Best-fit model VT+F+R10) using 1,000 ultrafast bootstrap (56–58). The M proteins identified for M5- and M6-like FH-binding patterns were selected and used for phylogenetic analysis using the aforementioned criterion, except the best fit model for substitution used was VT+F+G4. FigTree v1.4.4 was used to generate phylogenetic figures.

A collection of global *S. pyogenes* isolates composed of 148 M types were searched for FbaA sequences (32). These sequences were aligned with ClustalW (59) and the conservation of FH-binding amino acids displayed with ESpript 3.0 (60). The M proteins of strains positive for FbaA were used for phylogenetic analysis as above, except the best fit model for substitution was VT+F+I+G4.

## Acknowledgements

We thank Victor Nizet, Samira Dahesh, Elisabet Bjånes, and Jan-Willem Veening for contributions to the project. This work was supported by NIH R01AI154149 (PG) This research used resources of the Advanced Photon Source, a U.S. Department of Energy (DOE) Office of Science user facility operated for the DOE Office of Science by Argonne National Laboratory under Contract No. DE-AC02-06CH11357. Use of the Stanford Synchrotron Radiation Lightsource, SLAC National Accelerator Laboratory, is supported by the U.S. Department of Energy, Office of Science, Office of Basic Energy Sciences under Contract No. DE-AC02-76SF00515. The SSRL Structural Molecular Biology Program is supported by the DOE Office of Biological and Environmental Research, and by the National Institutes of Health, National Institute of General Medical Sciences (including P41GM103393). The contents of this publication are solely the responsibility of the authors and do not necessarily represent the official views of NIGMS or NIH.

## Author Contributions

Conceptualization: PG

Investigation: AK and KW

Formal Analysis: AK

Visualization: AK

Funding acquisition: PG

Project Administration: PG

Supervision: PG

Writing – original draft: AK and PG

Writing – review & editing: AK, KCW, and PG

## Supplementary Information

Supplementary Figures S1-S13.

Supplementary Tables S1-S3.

## References

1. Carapetis, J. R., Steer, A. C., Mulholland, E. K., and Weber, M. (2005) The global burden of group A streptococcal diseases Lancet Infect Dis 5, 685–694

2. Ghosh, P. (2018) Variation, Indispensability, and Masking in the M protein Trends Microbiol 26, 132–144

3. Whitnack, E., and Beachey, E. H. (1982) Antiopsonic activity of fibrinogen bound to M protein on the surface of group A streptococci J Clin Invest 69, 1042–1045

4. Macheboeuf, P., Buffalo, C., Fu, C. Y., Zinkernagel, A. S., Cole, J. N., Johnson, J. E. et al. (2011) Streptococcal M1 protein constructs a pathological host fibrinogen network Nature 472, 64–68

5. Thern, A., Stenberg, L., Dahlback, B., and Lindahl, G. (1995) Ig-binding surface proteins of Streptococcus pyogenes also bind human C4b-binding protein (C4BP), a regulatory component of the complement system J Immunol 154, 375–386

6. Buffalo, C. Z., Bahn-Suh, A. J., Hirakis, S. P., Biswas, T., Amaro, R. E., Nizet, V., et al. (2016) Conserved patterns hidden within group A Streptococcus M protein hypervariability recognize human C4b-binding protein Nat Microbiol 1, 16155

7. Kolesinski, P., McGowan, M., Botteaux, A., Smeesters, P. R., and Ghosh, P. (2024) Conservation of C4BP-binding sequence patterns in Streptococcus pyogenes M and Enn proteins J Biol Chem 300, 107478

8. Horstmann, R. D., Sievertsen, H. J., Knobloch, J., and Fischetti, V. A. (1988) Antiphagocytic activity of streptococcal M protein: selective binding of complement control protein factor H Proc Natl Acad Sci U S A 85, 1657–1661

9. Hong, K., Kinoshita, T., Takeda, J., Kozono, H., Pramoonjago, P., Kim, Y. U. et al. (1990) Inhibition of the alternative C3 convertase and classical C5 convertase of complement by group A streptococcal M protein Infect Immun 58, 2535–2541

10. Johnsson, E., Berggard, K., Kotarsky, H., Hellwage, J., Zipfel, P. F., Sjobring, U. et al. (1998) Role of the hypervariable region in streptococcal M proteins: binding of a human complement inhibitor J Immunol 161, 4894–4901

11. Gustafsson, M. C., Lannergard, J., Nilsson, O. R., Kristensen, B. M., Olsen, J. E., Harris, C. L., et al. (2013) Factor H binds to the hypervariable region of many Streptococcus pyogenes M proteins but does not promote phagocytosis resistance or acute virulence PLoS Pathog 9, e1003323

12. Swanson, J., Hsu, K. C., and Gotschlich, E. C. (1969) Electron microscopic studies on streptococci. I. M antigen J Exp Med 130, 1063–1091

13. Phillips, G. N., Jr., Flicker, P. F., Cohen, C., Manjula, B. N., and Fischetti, V. A. (1981) Streptococcal M protein: alpha-helical coiled-coil structure and arrangement on the cell surface Proc Natl Acad Sci U S A 78, 4689–4693

14. McNamara, C., Zinkernagel, A. S., Macheboeuf, P., Cunningham, M. W., Nizet, V., and Ghosh, P. (2008) Coiled-coil irregularities and instabilities in group A *Streptococcus* M1 are required for virulence Science 319, 1405–1408

15. Efstratiou, A., and Lamagni, T. (2022) Epidemiology of Streptococcus pyogenes In Streptococcus pyogenes: Basic Biology to Clinical Manifestations, 2nd Ed. Ferretti JJ, Stevens DL, and Fischetti VA, eds. Oklahoma City (OK)

16. Kotarsky, H., Hellwage, J., Johnsson, E., Skerka, C., Svensson, H. G., Lindahl, G. et al. (1998) Identification of a domain in human factor H and factor H-like protein-1 required for the interaction with streptococcal M proteins J Immunol 160, 3349–3354

17. Blackmore, T. K., Fischetti, V. A., Sadlon, T. A., Ward, H. M., and Gordon, D. L. (1998) M protein of the group A Streptococcus binds to the seventh short consensus repeat of human complement factor H Infect Immun 66, 1427–1431

18. Blackmore, T. K., Sadlon, T. A., Ward, H. M., Lublin, D. M., and Gordon, D. L. (1996) Identification of a heparin binding domain in the seventh short consensus repeat of complement factor H J Immunol 157, 5422–5427

19. Prosser, B. E., Johnson, S., Roversi, P., Herbert, A. P., Blaum, B. S., Tyrrell, J. et al. (2007) Structural basis for complement factor H linked age-related macular degeneration J Exp Med 204, 2277–2283

20. Schneider, M. C., Prosser, B. E., Caesar, J. J., Kugelberg, E., Li, S., Zhang, Q. et al. (2009) Neisseria meningitidis recruits factor H using protein mimicry of host carbohydrates Nature 458, 890–893

21. Marcinkiewicz, A. L., Brangulis, K., Dupuis, A. P., 2nd, Hart, T. M., Zamba-Campero, M., Nowak, T. A., et al. (2023) Structural evolution of an immune evasion determinant shapes pathogen host tropism Proc Natl Acad Sci U S A 120, e2301549120

22. Sharma, A. K., and Pangburn, M. K. (1996) Identification of three physically and functionally distinct binding sites for C3b in human complement factor H by deletion mutagenesis Proc Natl Acad Sci U S A 93, 10996–11001

23. Pandiripally, V., Gregory, E., and Cue, D. (2002) Acquisition of regulators of complement activation by Streptococcus pyogenes serotype M1 Infect Immun 70, 6206–6214

24. Terao, Y., Kawabata, S., Kunitomo, E., Murakami, J., Nakagawa, I., and Hamada, S. (2001) Fba, a novel fibronectin-binding protein from Streptococcus pyogenes, promotes bacterial entry into epithelial cells, and the fba gene is positively transcribed under the Mga regulator Mol Microbiol 42, 75–86

25. Perez-Caballero, D., Garcia-Laorden, I., Cortes, G., Wessels, M. R., de Cordoba, S. R., and Alberti, S. (2004) Interaction between complement regulators and Streptococcus pyogenes: binding of C4b-binding protein and factor H/factor H-like protein 1 to M18 strains involves two different cell surface molecules J Immunol 173, 6899–6904

26. Agrahari, G., Liang, Z., Mayfield, J. A., Balsara, R. D., Ploplis, V. A., and Castellino, F. J. (2013) Complement-mediated opsonization of invasive group A Streptococcus pyogenes strain AP53 is regulated by the bacterial two-component cluster of virulence responder/sensor (CovRS) system J Biol Chem 288, 27494–27504

27. Pandiripally, V., Wei, L., Skerka, C., Zipfel, P. F., and Cue, D. (2003) Recruitment of complement factor H-like protein 1 promotes intracellular invasion by group A streptococci Infect Immun 71, 7119–7128

28. Kumar, P., and Woolfson, D. N. (2021) Socket2: A Program for Locating, Visualising, and Analysing Coiled-coil Interfaces in Protein Structures Bioinformatics 37, 4575–4577

29. Strelkov, S. V., and Burkhard, P. (2002) Analysis of alpha-helical coiled coils with the program TWISTER reveals a structural mechanism for stutter compensation J Struct Biol 137, 54–64

30. Lawrence, M. C., and Colman, P. M. (1993) Shape complementarity at protein/protein interfaces J Mol Biol 234, 946–950

31. Sanderson-Smith, M., De Oliveira, D. M., Guglielmini, J., McMillan, D. J., Vu, T., Holien, J. K. et al. (2014) A systematic and functional classification of *Streptococcus pyogenes* that serves as a new tool for molecular typing and vaccine development J Infect Dis 210, 1325–1338

32. Frost, H. R., Davies, M. R., Delforge, V., Lakhloufi, D., Sanderson-Smith, M., Srinivasan, V. et al. Analysis of Global Collection of Group A Streptococcus Genomes Reveals that the Majority Encode a Trio of M and M-Like Proteins mSphere

33. Frost, H. R., Guglielmini, J., Duchene, S., Lacey, J. A., Sanderson-Smith, M., Steer, A. C. et al. Promiscuous evolution of Group A Streptococcal M and M-like proteins Microbiology (Reading)

34. Abramson, J., Adler, J., Dunger, J., Evans, R., Green, T., Pritzel, A. et al. (2024) Accurate structure prediction of biomolecular interactions with AlphaFold 3 Nature 630, 493–500

35. Haapasalo, K., Jarva, H., Siljander, T., Tewodros, W., Vuopio-Varkila, J., and Jokiranta, T. S. (2008) Complement factor H allotype 402H is associated with increased C3b opsonization and phagocytosis of Streptococcus pyogenes Mol Microbiol 70, 583–594

36. Yu, J., Wiita, P., Kawaguchi, R., Honda, J., Jorgensen, A., Zhang, K. et al. (2007) Biochemical analysis of a common human polymorphism associated with age-related macular degeneration Biochemistry 46, 8451–8461

37. Giannakis, E., Jokiranta, T. S., Male, D. A., Ranganathan, S., Ormsby, R. J., Fischetti, V. A. et al. (2003) A common site within factor H SCR 7 responsible for binding heparin, C-reactive protein and streptococcal M protein Eur J Immunol 33, 962–969

38. Perez-Caballero, D., Alberti, S., Vivanco, F., Sanchez-Corral, P., and Rodriguez de Cordoba, S. (2000) Assessment of the interaction of human complement regulatory proteins with group A Streptococcus. Identification of a high-affinity group A Streptococcus binding site in FHL-1 Eur J Immunol 30, 1243–1253

39. Haapasalo, K., Vuopio, J., Syrjanen, J., Suvilehto, J., Massinen, S., Karppelin, M. et al. (2012) Acquisition of complement factor H is important for pathogenesis of Streptococcus pyogenes infections: evidence from bacterial in vitro survival and human genetic association J Immunol 188, 426–435

40. Schwarz-Linek, U., Werner, J. M., Pickford, A. R., Gurusiddappa, S., Kim, J. H., Pilka, E. S. et al. (2003) Pathogenic bacteria attach to human fibronectin through a tandem beta-zipper Nature 423, 177–181

41. Persson, J., Beall, B., Linse, S., and Lindahl, G. (2006) Extreme sequence divergence but conserved ligand-binding specificity in *Streptococcus pyogenes* M protein PLoS Pathog 2, e47

42. Berggard, K., Johnsson, E., Morfeldt, E., Persson, J., Stalhammar-Carlemalm, M., and Lindahl, G. (2001) Binding of human C4BP to the hypervariable region of M protein: a molecular mechanism of phagocytosis resistance in Streptococcus pyogenes Mol Microbiol 42, 539–551

43. Carlsson, F., Berggard, K., Stalhammar-Carlemalm, M., and Lindahl, G. (2003) Evasion of phagocytosis through cooperation between two ligand-binding regions in *Streptococcus pyogenes* M protein J Exp Med 198, 1057–1068

44. Sandin, C., Carlsson, F., and Lindahl, G. (2006) Binding of human plasma proteins to *Streptococcus pyogenes* M protein determines the location of opsonic and non-opsonic epitopes Mol Microbiol 59, 20–30

45. Kotarsky, H., Gustafsson, M., Svensson, H. G., Zipfel, P. F., Truedsson, L., and Sjobring, U. (2001) Group A streptococcal phagocytosis resistance is independent of complement factor H and factor H-like protein 1 binding Mol Microbiol 41, 817–826

46. Otwinowski, Z., and Minor, W. (1997) Processing of X-ray diffraction data collected in oscillation mode Method Enzymol 276, 307–326

47. Karplus, P. A., and Diederichs, K. (2012) Linking crystallographic model and data quality Science 336, 1030–1033

48. Potterton, L., Agirre, J., Ballard, C., Cowtan, K., Dodson, E., Evans, P. R. et al. (2018) CCP4i2: the new graphical user interface to the CCP4 program suite Acta Crystallogr D Struct Biol 74, 68–84

49. Liebschner, D., Afonine, P. V., Baker, M. L., Bunkóczi, G., Chen, V. B., Croll, T. I. et al. (2019) Macromolecular structure determination using X-rays, neutrons and electrons: recent developments in *Acta* Crystallogr D 75, 861–877

50. McCoy, A. J., Grosse-Kunstleve, R. W., Adams, P. D., Winn, M. D., Storoni, L. C., and Read, R. J. (2007) *Phaser* crystallographic software Journal of Applied Crystallography 40, 658–674

51. Mirdita, M., Schutze, K., Moriwaki, Y., Heo, L., Ovchinnikov, S., and Steinegger, M. (2022) ColabFold: making protein folding accessible to all Nat Methods 19, 679–682

52. Afonine, P. V., Grosse-Kunstleve, R. W., Echols, N., Headd, J. J., Moriarty, N. W., Mustyakimov, M. et al. (2012) Towards automated crystallographic structure refinement with phenix.refine Acta Crystallographica Section D 68, 352–367

53. Krissinel, E., and Henrick, K. (2007) Inference of macromolecular assemblies from crystalline state J Mol Biol 372, 774–797

54. Edgar, R. C. (2004) MUSCLE: multiple sequence alignment with high accuracy and high throughput Nucleic Acids Res 32, 1792–1797

55. Tamura, K., Stecher, G., and Kumar, S. (2021) MEGA11: Molecular Evolutionary Genetics Analysis Version 11 Mol Biol Evol 38, 3022–3027

56. Minh, B. Q., Schmidt, H. A., Chernomor, O., Schrempf, D., Woodhams, M. D., von Haeseler, A. et al. (2020) IQ-TREE 2: New Models and Efficient Methods for Phylogenetic Inference in the Genomic Era Mol Biol Evol 37, 1530–1534

57. Kalyaanamoorthy, S., Minh, B. Q., Wong, T. K. F., von Haeseler, A., and Jermiin, L. S. (2017) ModelFinder: fast model selection for accurate phylogenetic estimates Nat Methods 14, 587–589

58. Hoang, D. T., Chernomor, O., von Haeseler, A., Minh, B. Q., and Vinh, L. S. (2018) UFBoot2: Improving the Ultrafast Bootstrap Approximation Mol Biol Evol 35, 518–522

59. Larkin, M. A., Blackshields, G., Brown, N. P., Chenna, R., McGettigan, P. A., McWilliam, H. et al. (2007) Clustal W and Clustal X version 2.0 Bioinformatics 23, 2947–2948

60. Robert, X., and Gouet, P. (2014) Deciphering key features in protein structures with the new ENDscript server Nucleic Acids Res 42, W320–324

