## Supplemental Figures and Tables for "Structural mechanisms for the recruitment of factor H by *Streptococcus pyogenes*"

**Supplemental Figures S1-S13**

**Supplemental Tables S1-S3**

Supplemental Figure S1

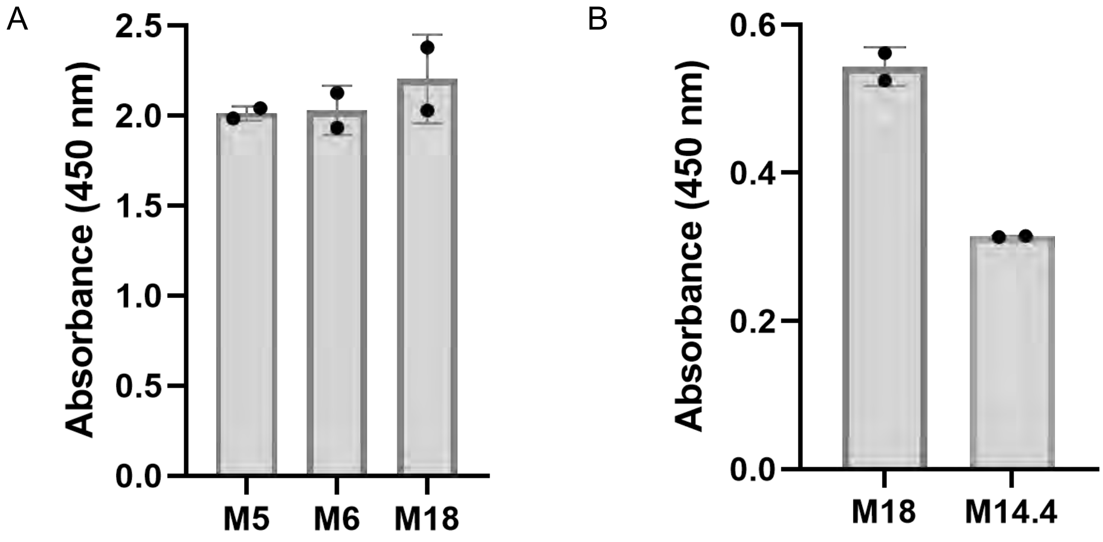

**Figure S1. FH binding to M proteins.**

**A.** Binding of soluble, intact FH to immobilized intact $His_6$-M5, $His_6$-M6, and $His_6$-M18 proteins, as evaluated by ELISA. Bound FH was detected with an anti-FH monoclonal antibody. Data from two biological replicates are presented with means and standard deviations.

**B.** Same as panel A, but for $His_6$-M18 and $His_6$-M14.4 proteins.

Supplemental Figure S2

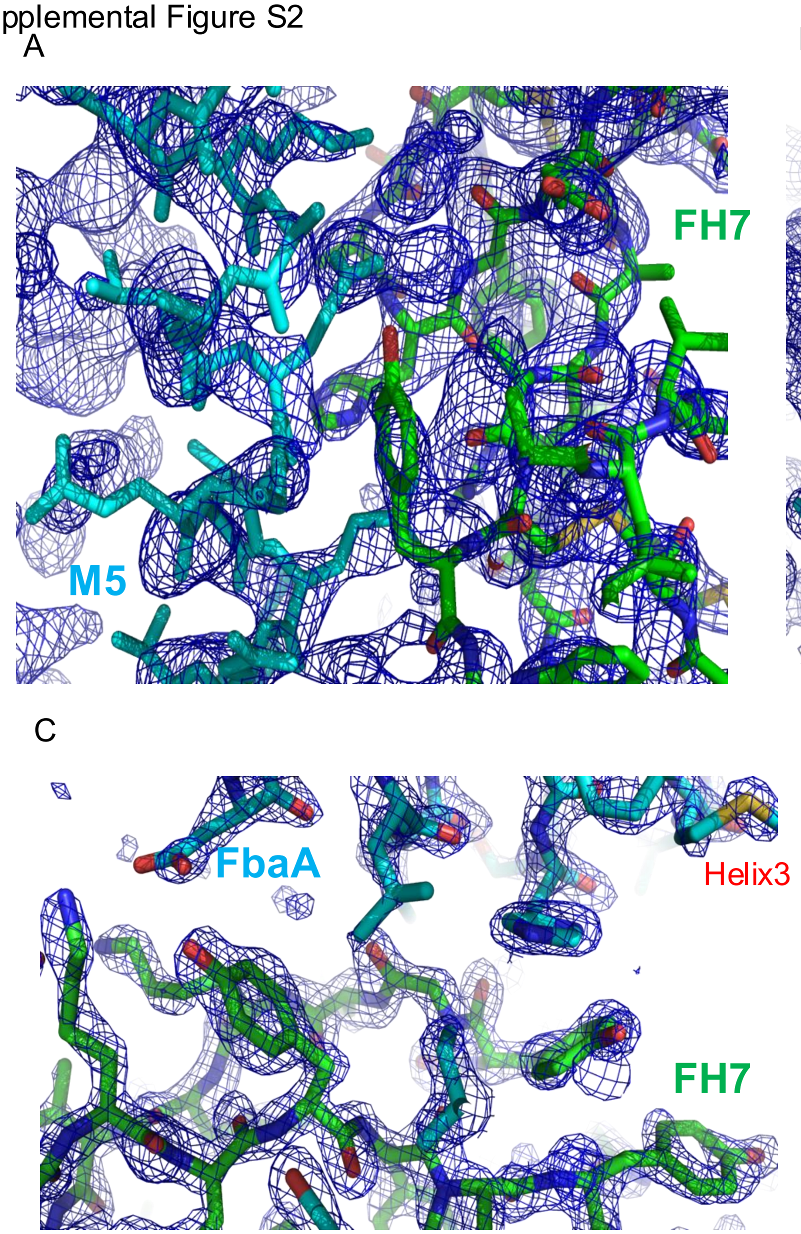

**Figures S2. Electron Density.** Composite omit 2mFo-DFc electron density map contoured at 2σ for the (**A**) M5/FH(6-7), (**B**) M6/FH(6-7), and (**C**) FbaA/FH(6-7) complexes. M5 protein, M6 protein, and FbaA are in cyan, and FH in green.

Supplemental Figure S3

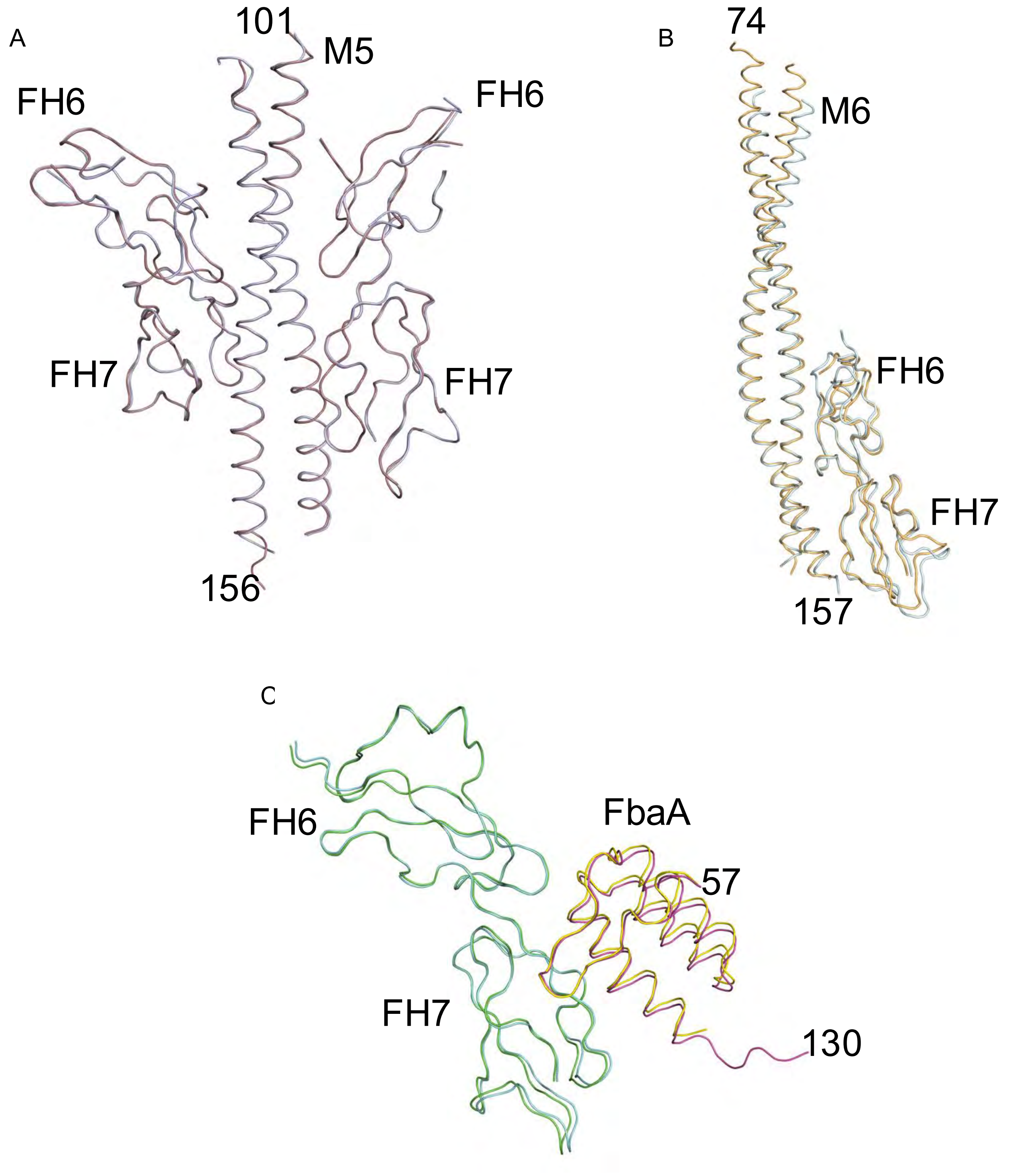

**Figure S3. M protein-FH complexes in the asymmetric unit.**
Superposition of the two (**A**) M5/FH(6-7), (**B**) M6/FH(6-7), and (**C)** FbaA/FH(6-7) complexes in the asymmetric unit of each respective crystal.

Supplemental Figure S4

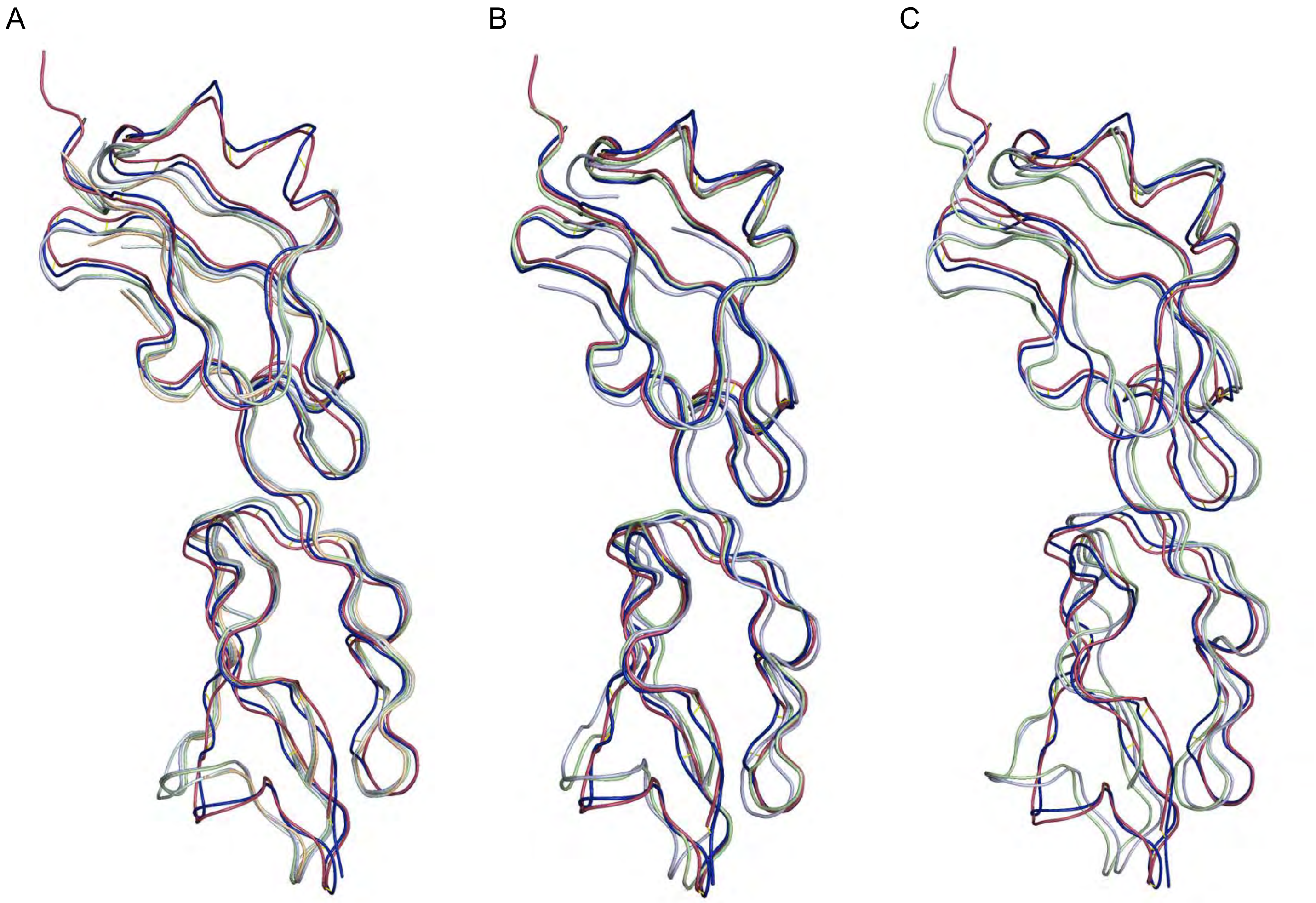

**Figure S4. Stable conformation of FH(6-7).** Superposition of FH(6-7) when bound to *N. meningitidis* fHbp (red, PDB:2w80), *B. burgdorferi* CspZ (blue, PDB: 7ZJM), and (**A**) M5 protein (green, cyan, light blue and wheat), (**B**) M6 protein (green and light blue), and (**C**) FbaA (green and light blue).

Supplemental Figure S5

A

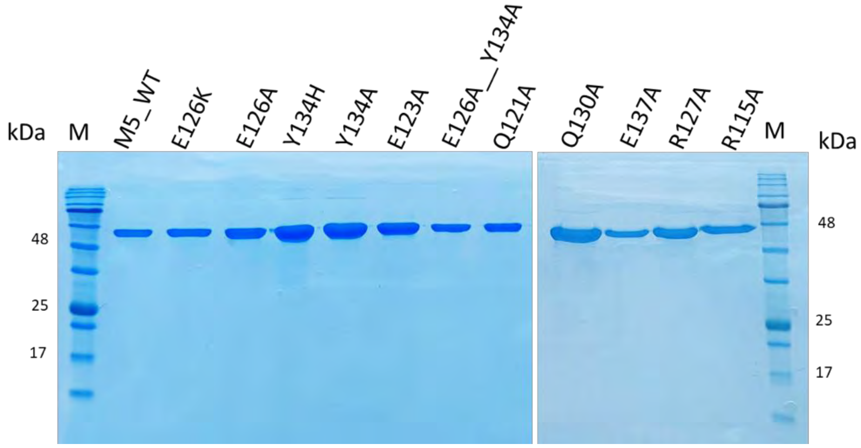

B

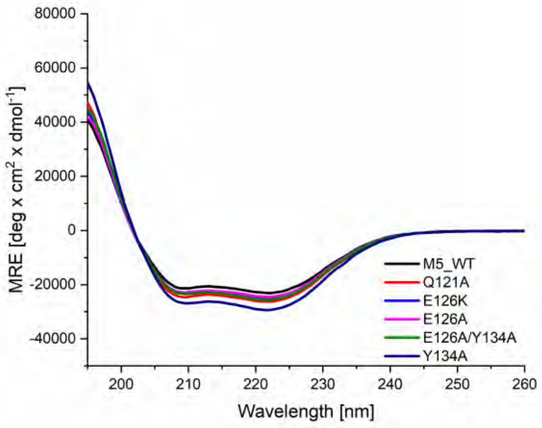

C

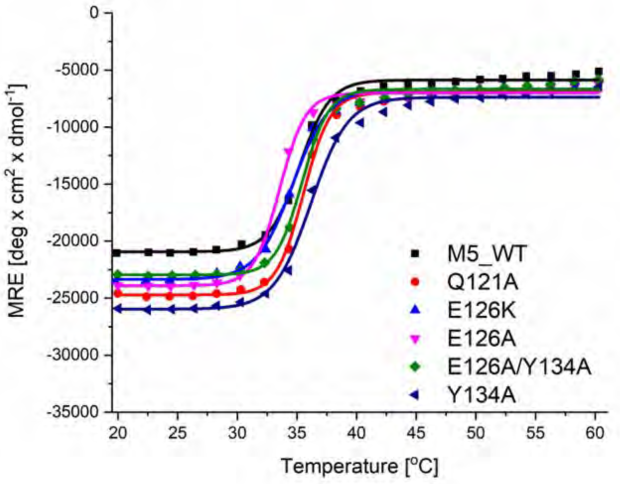

D

| Name | Tm (°C) |
| --- | --- |
| M5_WT | 35.85 ± 0.91 |
| Q121A | 34.66 ± 1.22 |
| E126K | 34.86 ± 0.31 |
| E126A | 34.79 ± 1.76 |
| E126A/Y134A | 35.74 ± 0.45 |
| Y134A | 36.17 ± 0 |

E

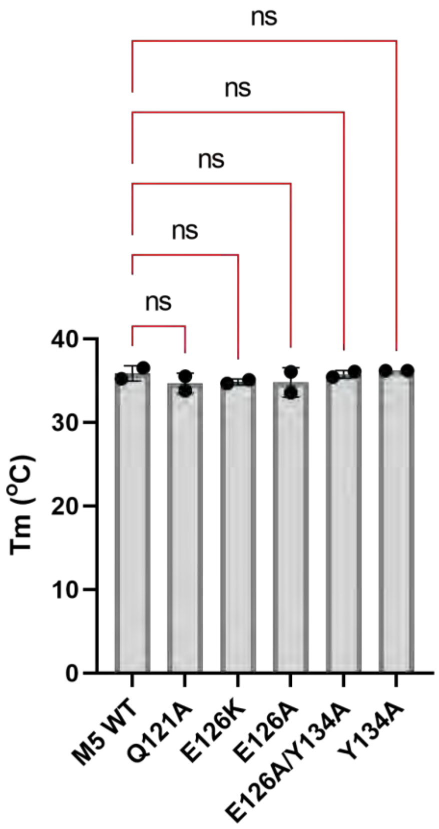

**Figure S5. Structure and stability of wild-type and mutant His$_6$-M5 proteins.**
**A.** SDS-PAGE of purified intact wild-type and mutant His$_6$-M5 proteins.
**B.** CD spectra of intact wild-type and mutant M5 proteins at 25 °C. The data are an average of two independent measurements.
**C.** Thermal melting curves of intact wild-type and mutant M5 proteins, as monitored by the CD signal at 222 nm. The data are an average of two independent measurements.
**D.** Tabulation of Tm's determined from panel C.
**E.** Statistical analysis of Tm's, determined by one-way ANOVA. P-value for ns > 0.123.

Supplemental Figure S6

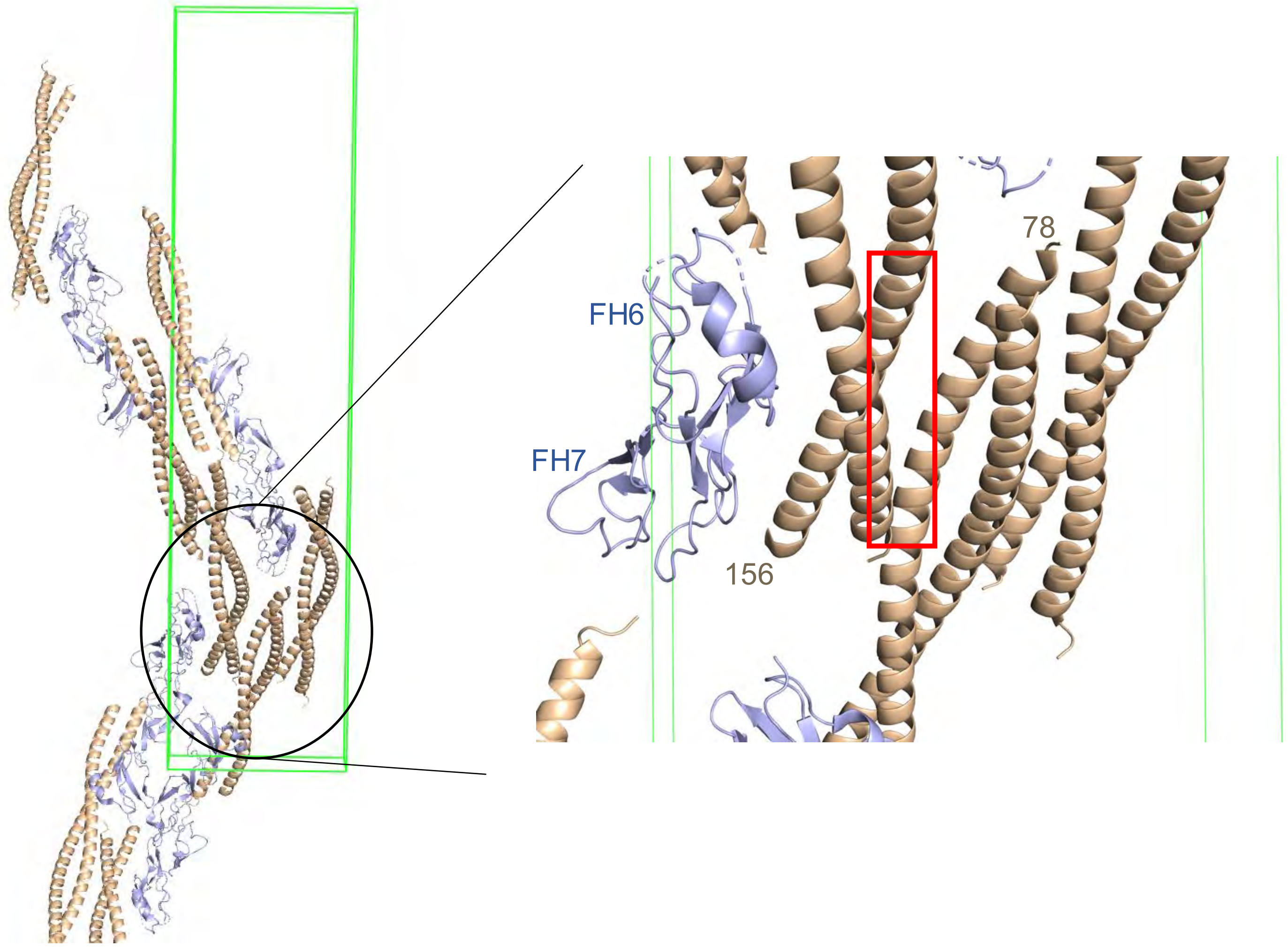

**Figure S6. M6/FH(6-7) crystal packing.** The green box indicates the unit cell, with M6 protein in wheat and FH(6-7) in light blue. The red box indicates an FH-binding site in M6 protein that is unoccupied and instead forms a contact with a crystallographically related M6 protein dimer.

Supplemental Figure S7

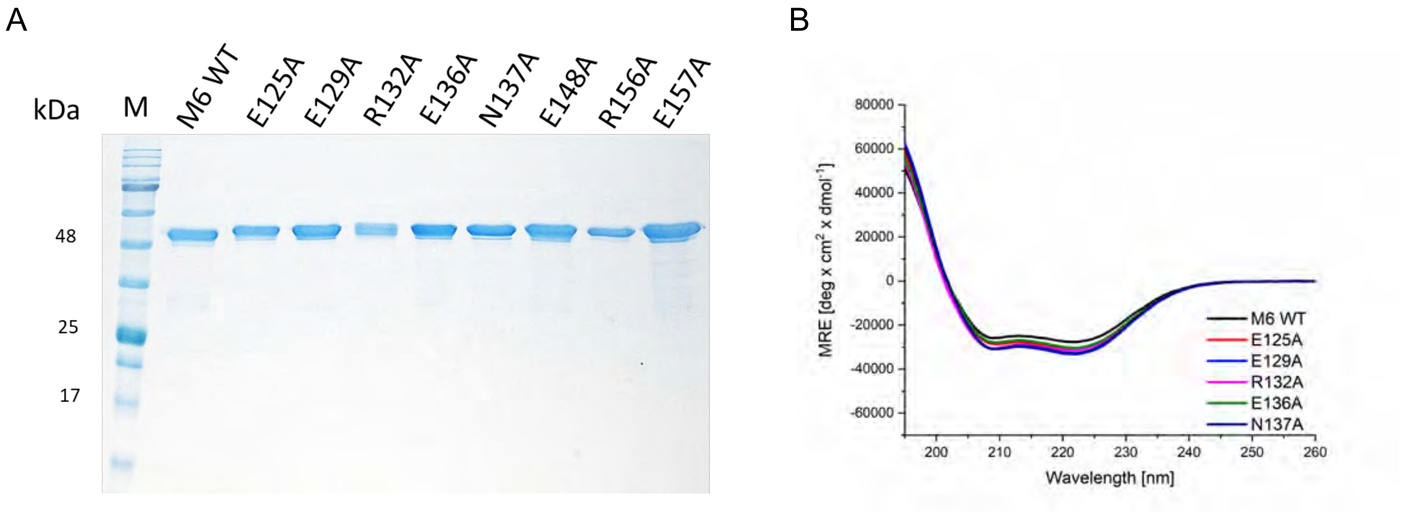

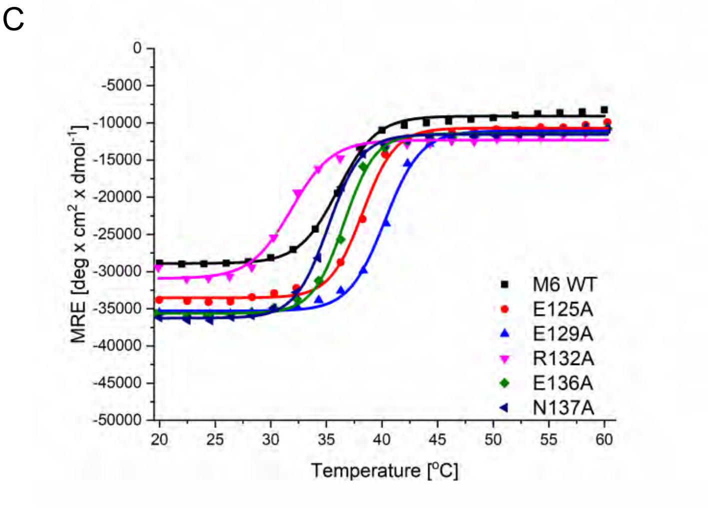

D

| Name | Tm ($^o$C) |
|---|---|
| **M6 WT** | 36.12 ± 0.11 |
| **E125A** | 38.74 ± 0.79 |
| **E129A** | 39.78 ± 0.65 |
| **R132A** | 32.03 ± 0.28 |
| **E136A** | 37.43 ± 1.23 |
| **N137A** | 34.98 ± 0.25 |

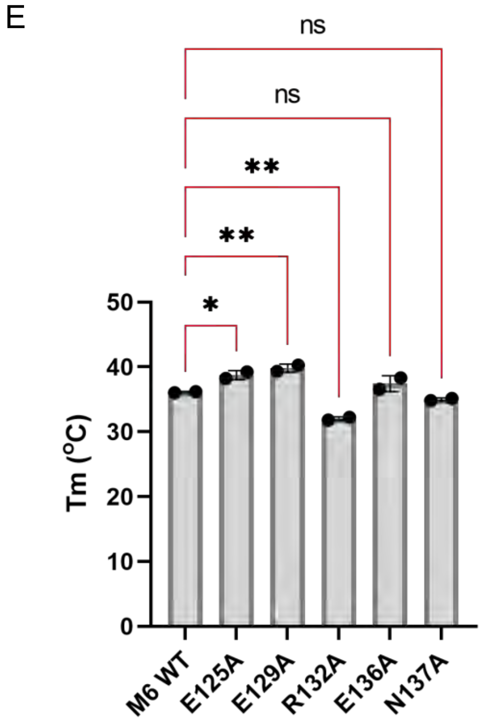

**Figure S7. Structure and stability of WT and mutant His$_6$-M6 proteins.**

**A.** SDS-PAGE of purified intact wild-type and mutant His$_6$-M6 proteins.

**B.** CD spectra of intact wild-type and mutant M6 proteins at 25 °C. The data are an average of two independent measurements.

**C.** Thermal melting curves of intact wild-type and mutant M6 proteins, as monitored by the CD signal at 222 nm. The data are an average of two independent measurements.

**D.** Tabulation of Tm's determined from panel C.

**E.** Statistical analysis of Tm's determined by one-way ANOVA. p-values are as follows: ns > 0.123, * ≤ 0.033, and ** ≤ 0.0021.

Supplemental Figure S8

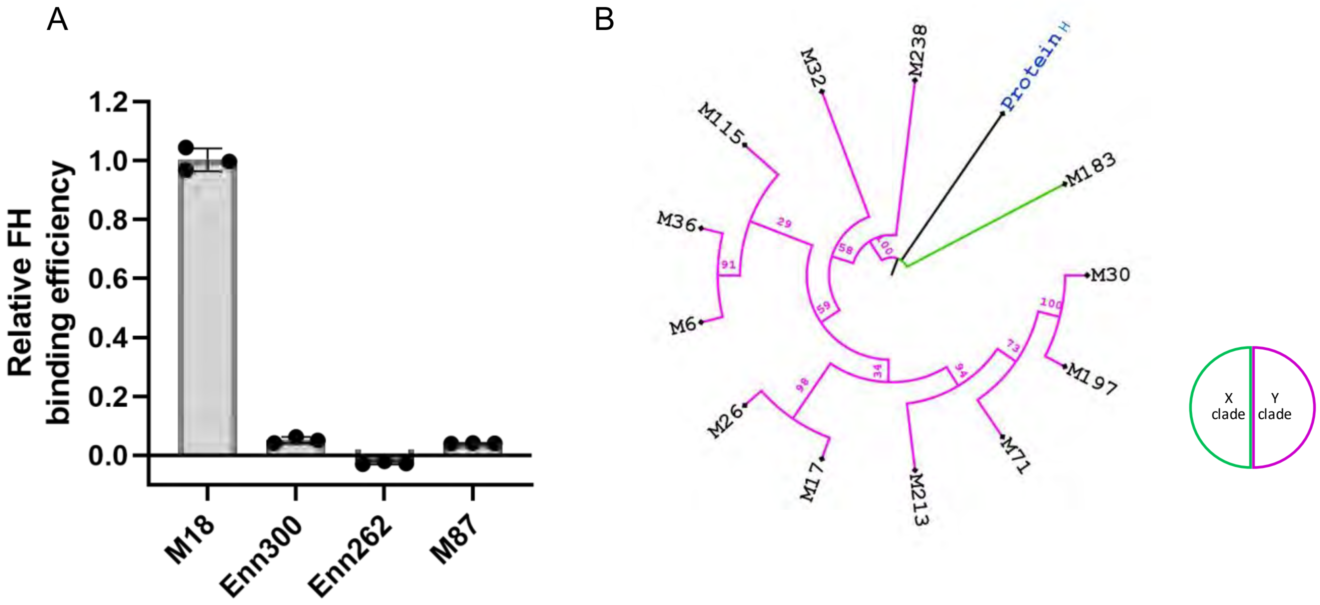

**Figure S8. Threshold for M6 FH-binding pattern and phylogenetics.**
**A.** Binding of soluble intact FH to immobilized intact $His_6$-M18, $His_6$-M87, $His_6$-Enn300, and $His_6$-Enn262 proteins, as evaluated by ELISA. Bound FH was detected with an anti-FH monoclonal antibody. Values were normalized by FH binding to $His_6$-M18 protein. Data from three biological replicates are presented with means and standard deviations.
**B.** Phylogenetic analysis of M proteins with M6 FH-binding pattern. Protein H was used as an outgroup.

Supplemental Figure S9

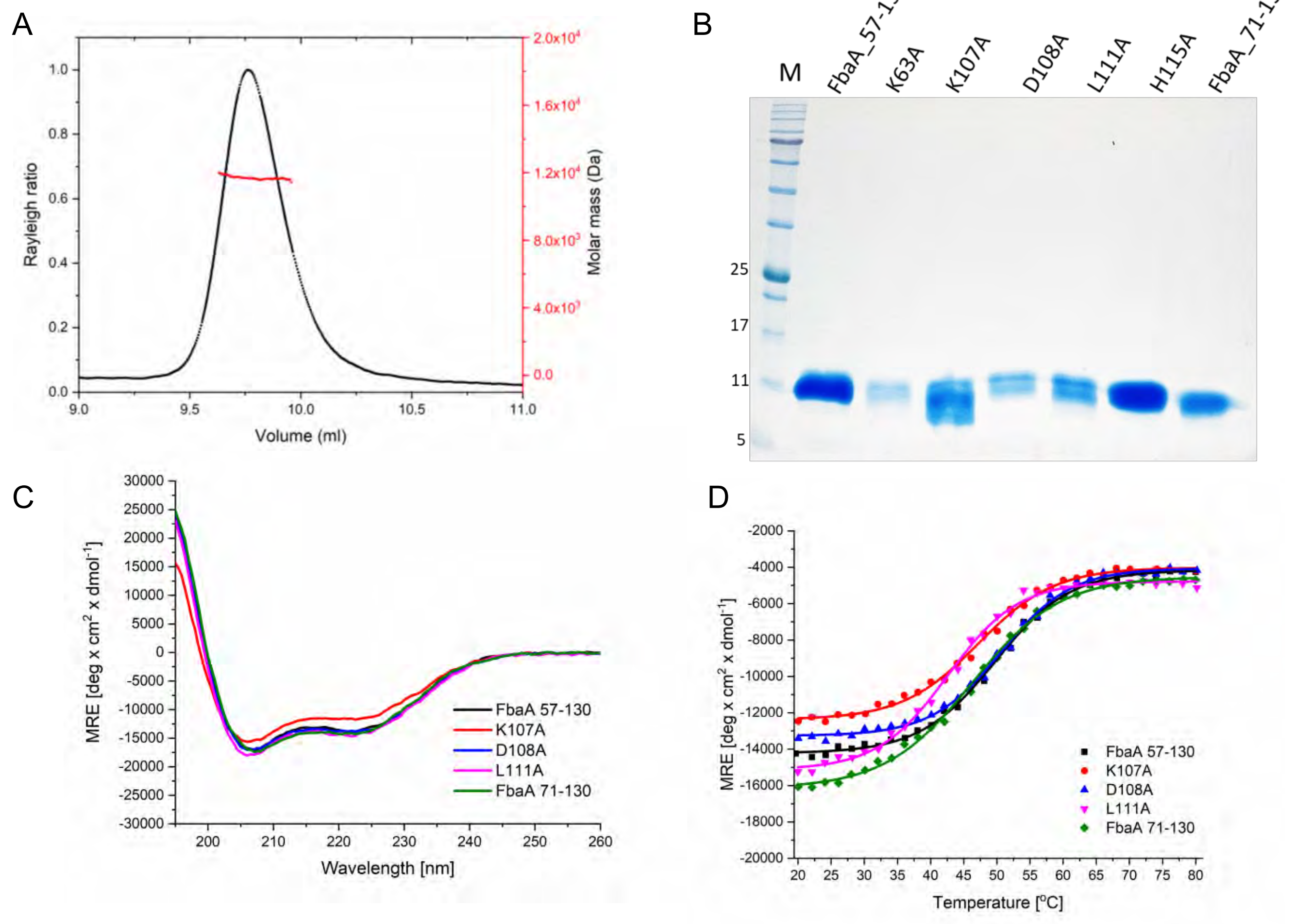

E

| Name | Tm (°C) |
|---|---|
| **FbaA 57-130** | 51.36 ± 2.31 |
| **K107A** | 48.92 ± 2.43 |
| **D108A** | 52.75 ± 3.00 |
| **L111A** | 44.00 ± 1.86 |
| **FbaA 71-130** | 48.00 ± 2.42 |

F

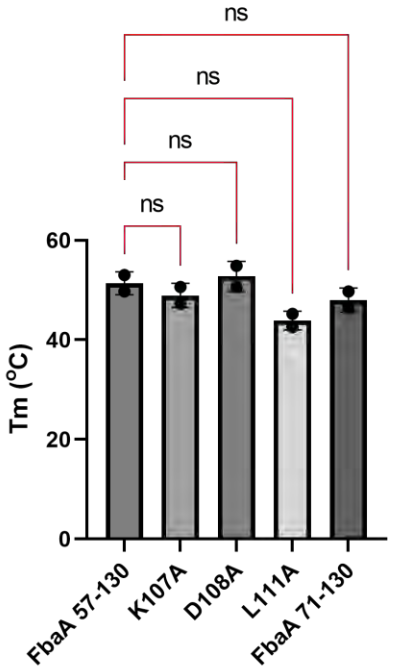

**Figure S9. Wild-type and mutant FbaA (57-130).**
**A.** Molecular mass of FbaA (57-130) determined by SEC-MALS.
**B.** SDS-PAGE of wild-type and mutant $His_6$-FbaA (57-130).
**C.** CD spectra of wild-type and mutant FbaA (57-130) at 25 °C. The data are an average of two independent measurements.
**D.** Thermal melting curves of intact wild-type and mutant FbaA (57-130), as monitored by the CD signal at 222 nm. The data are an average of two independent measurements.
**E.** Tabulation of Tm's determined from panel D.
**F.** Statistical analysis of Tm's, determined by ordinary one-way ANOVA. p-value for ns > 0.123.

Supplemental Figure S10

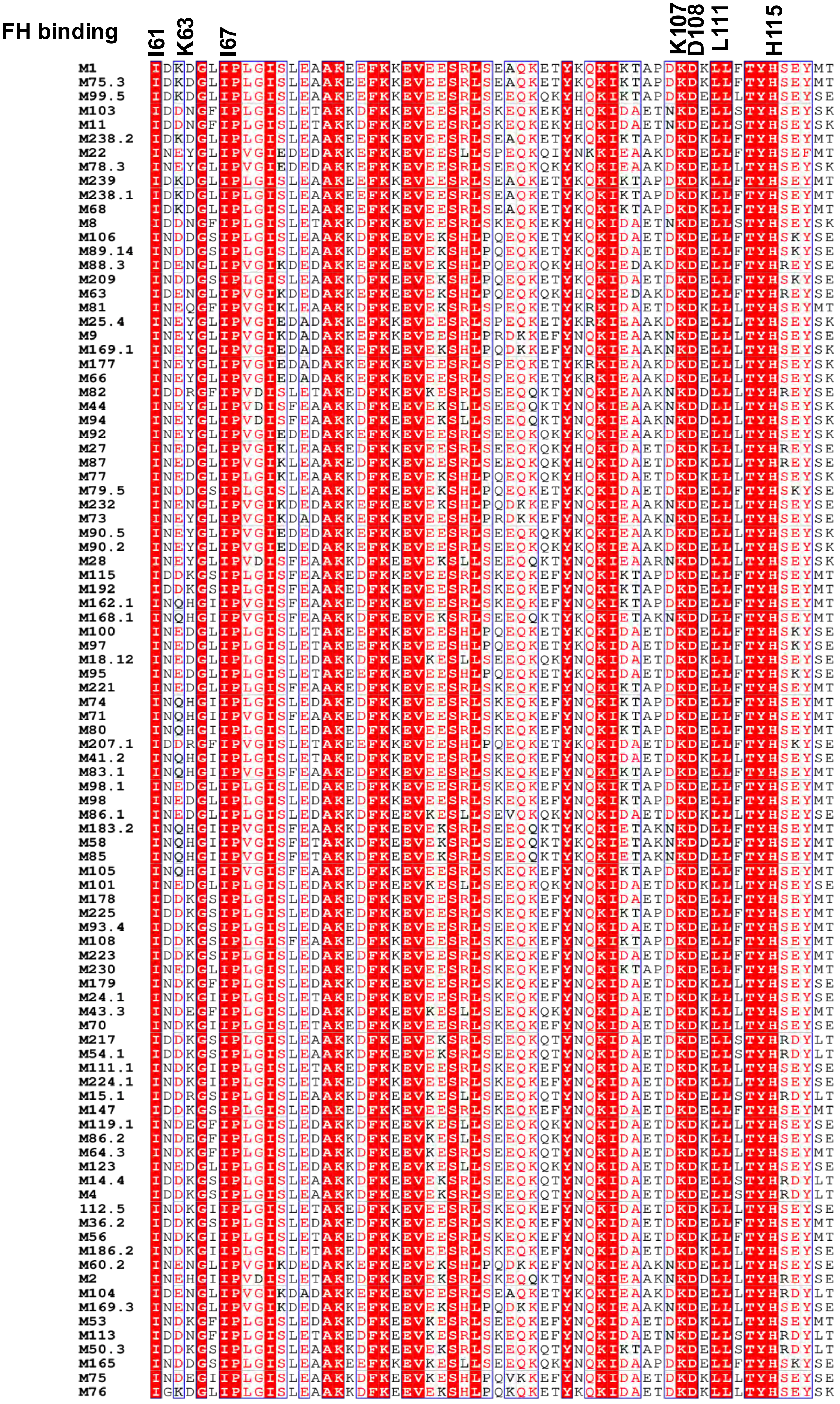

**Figure S10. FH-binding sequence in FbaA.**
Sequence alignment of the loop and three-helix bundle region from FbaA belonging to the M1 strain with FbaA from other M types. FH-interacting amino acids at the top.

Supplemental Figure S11

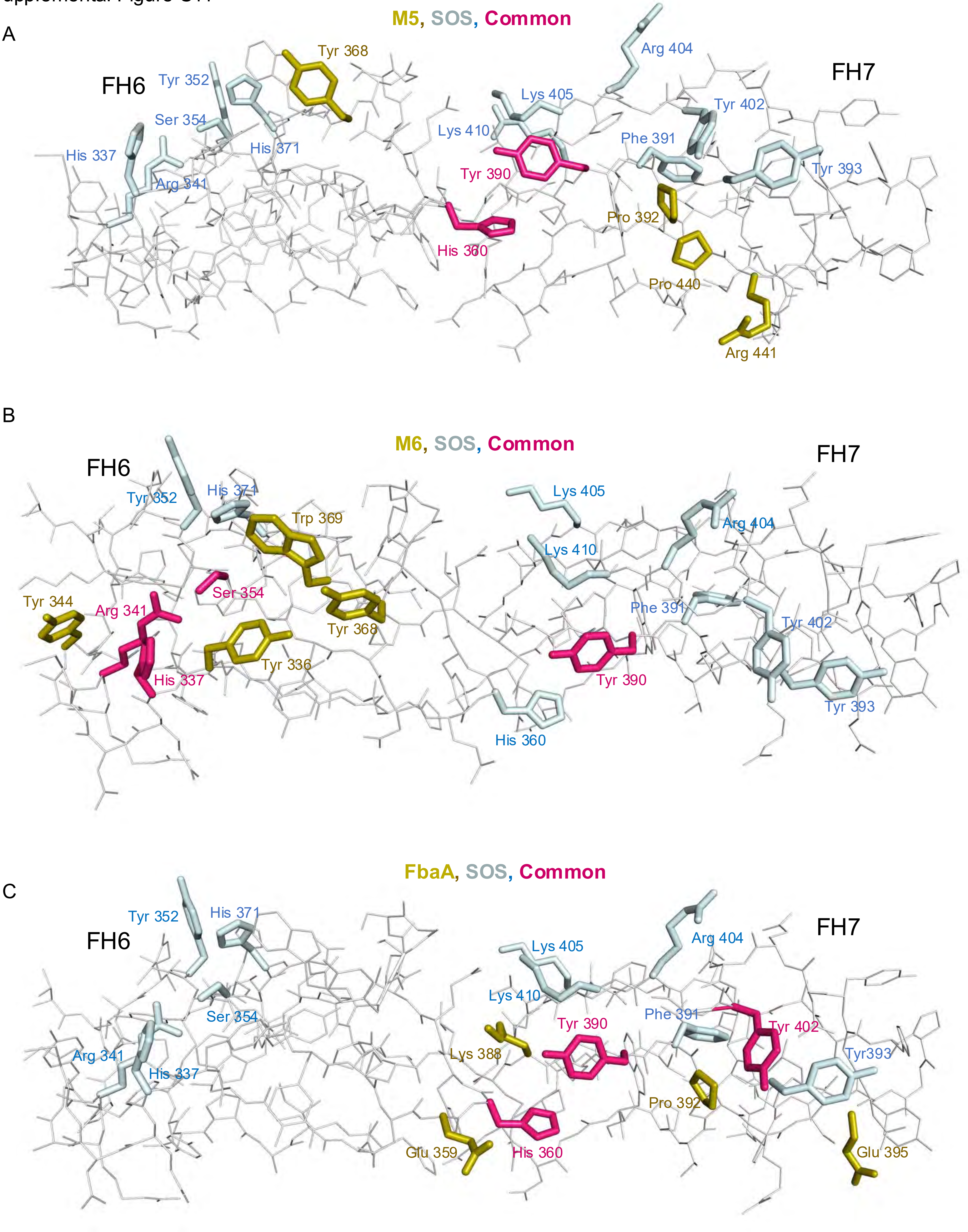

**Figure S11. Common FH amino acids for SOS.**
**A.** FH domains 6 and 7 with side chains that contact only M5 protein in gold, only SOS in pale blue, and those in common in red.
**B.** The same as panel A, but for M6 protein.
**C.** The same as panel A, but for FbaA.

Supplemental Figure S12

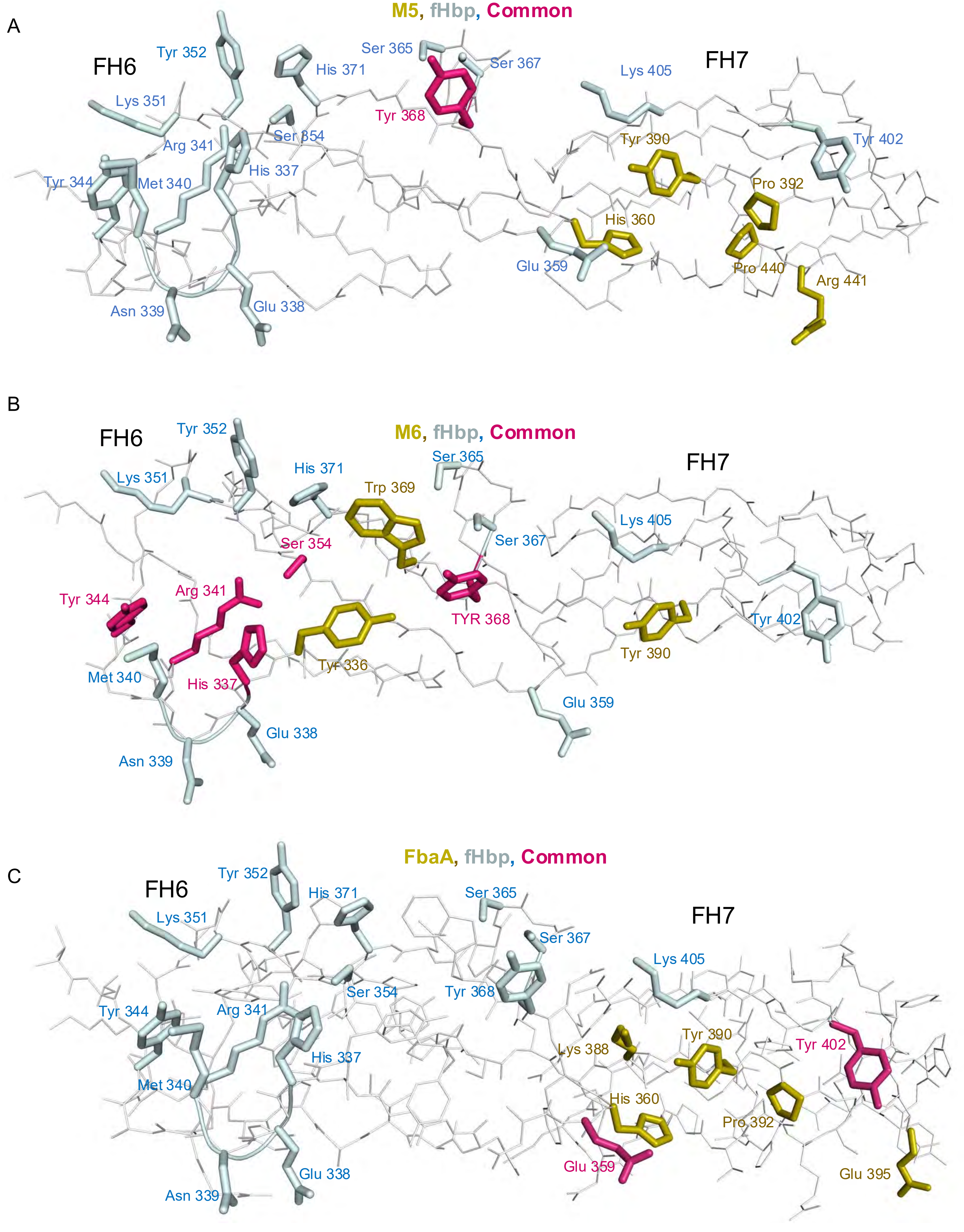

**Figure S12. Common FH amino acids for fHbp.**
**A.** FH domains 6 and 7 with side chains that contact only M5 protein in gold, only fHbp in pale blue, and those in common in red.
**B.** The same as panel A, but for M6 protein.
**C.** The same as panel A, but for FbaA.

Supplemental Figure S13

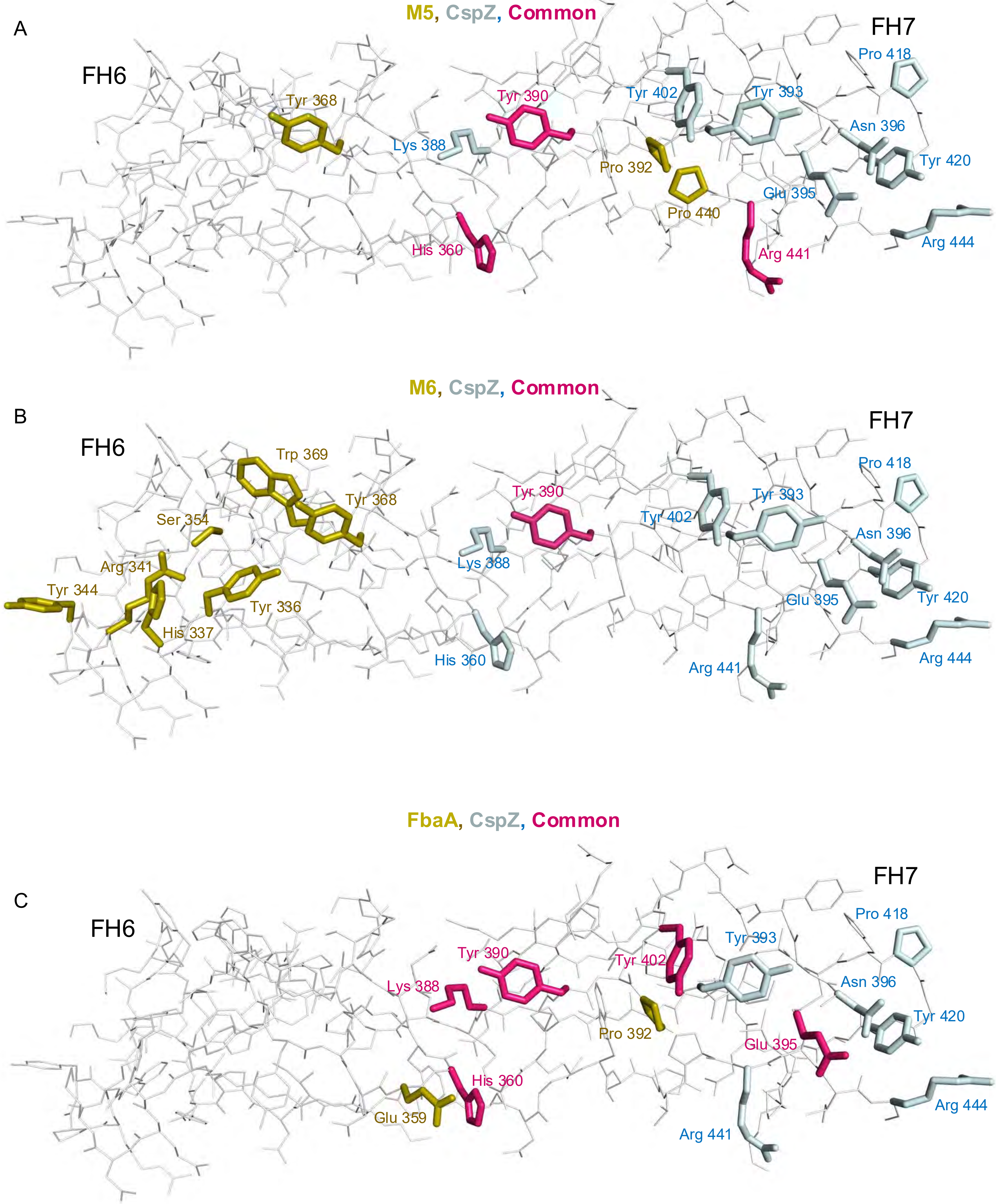

**Figure S13. Common FH amino acids for CspZ.**
**A.** FH domains 6 and 7 with side chains that contact only M5 protein in gold, only CspZ in pale blue, and those in common in red.
**B.** The same as panel A, but for M6 protein.
**C.** The same as panel A, but for FbaA.

**Table S1.** Crystallographic data collection and model refinement.

| | M5-FH | M6-FH | FbaA-FH |
|---|---|---|---|
| **Data collection** | | | |
| Wavelength (Å) | 1.0 | 1.0 | 1.54 |
| Resolution range (Å) | 50.00-2.25 (2.29-2.25) | 44.90-1.90 (1.94-1.90) | 36.19-1.82 (1.86-1.82) |
| Space group | $P\ 3_2$ | $P\ 2\ 2_1\ 2_1$ | $P\ 1\ 2_1\ 1$ |
| **Cell dimensions** | | | |
| a, b, c (Å) | 62.7, 62.7, 199.3 | 33.4, 69.5, 293.9 | 28.5, 79.0, 89.9 |
| α, β, γ (°) | 90.0, 90.0,120.0 | 90.0, 90.0, 90.0 | 90.0, 91.8, 90.0 |
| Total reflections | 266,494 | 711,806 | 69,389 |
| Unique reflections | 41,276 | 55,717 | 35,778 |
| Redundancy | 6.5 (5.6) | 12.8 (12.7) | 1.9 (1.9) |
| Completeness (%) | 99.5 (98.7) | 99.8 (99.9) | 99.8 (99.6) |
| I/σ(I) | 11.4 (0.5) | 9.8 (0.5) | 6.6 (1.6) |
| Rmeas | 0.23 (1.859) | 0.130 (5.955) | 0.094 (0.809) |
| $CC_{1/2}$ | 0.948 (0.070) | 0.999 (0.304) | 0.996 (0.543) |
| Wilson B-factor ($Å^2$) | 45.2 | 36.6 | 18.6 |
| **Refinement** | | | |
| Resolution range (Å) | 47.72 - 2.25 | 44.89 - 1.90 | 36.19 - 1.82 |
| No. of reflections (work/test set) | 41,213/4,081 | 55,577/5,535 | 35,742/3,568 |
| Rwork/Rfree | 0.253/0.287 | 0.260/0.302 | 0.226/0.275 |
| No. of non-hydrogen atoms for Protein | 4,544 | 4,075 | 3,016 |
| No. of non-hydrogen atoms for Ligands | None | 12 (ethylene glycol) | 48 (MES) |
| No. of non-hydrogen atoms (water) | 70 | 98 | 312 |
| L-test for twinning* | < \|L\| > = 0.41, < L2 > = 0.23 | < \|L\| > = 0.50, < L2 > = 0.33 | < \|L\| > = 0.50, < L2 > = 0.33 |
| **r.m.s. deviations** | | | |
| Bonds (Å) | 0.0025 | 0.0050 | 0.0040 |
| Angles (°) | 0.47 | 0.69 | 0.79 |
| **Ramachandran plot** | | | |
| Favored (%)/allowed (%) | 95.15/4.85 | 99.08/0.92 | 98.96/1.04 |
| Outliers (%) | 0 | 0 | 0 |
| Rotamer outliers (%) | 1.81 | 0 | 0 |
| Clashscore | 6.87 | 3.87 | 4.33 |
| Number of TLS groups | 0 | 23 | 10 |
| PDB code | 9MMU | 9MMX | 9MLU |

*Theoretical values of < |L| >, < L2 > for acentric reflections are 0.5, 0.333 respectively for untwined datasets, and 0.375, 0.2 for perfectly twinned datasets.

**Table S2. M5 FH-binding pattern in M, Enn and Mrp proteins.**
Red horizontal line indicates cut-off.

| M | Position | Sequence | Score |
|---|---|---|---|
| **29** | **121** | **QRETLEKNNDDLTYKNDDL** | **5.6** |
| **37** | **127** | **QKDKLEKEVQEKEYNNDGL** | **5.4** |
| **142** | **156** | **QKETLEREVQNTQYNNETL** | **5.4** |
| **5** | **121** | **QKETLEREVQNTQYNNETL** | **5.4** |
| **46** | **183** | **QKENLEKEVAEATYKNETL** | **5.2** |
| **54** | **115** | **QRQNLEKEVAETKYKNETL** | **5** |
| **M2_enn127** | **132** | **KLEAINKELNENYYKLQDG** | **5** |
| **95** | **253** | **RLNAQERMYEAFLYQAKDI** | **4.7** |
| **14** | **134** | **QKERLEKKVQETEYNNGEL** | **4.7** |
| **74** | **141** | **QKETLERQVQEKEHNNEAL** | **4.6** |
| **14.4** | **162** | **QKEKLEKQVQEKEHNNEAL** | **4.6** |
| **19** | **163** | **QKETLERQVQEAQHNNNEL** | **4.5** |
| **49** | **88** | **ELEERQKNLEKLEHQSQVA** | **4.4** |
| **151** | **81** | **ELEERQKNLEKLEHQFQVA** | **4.4** |
| **47** | **150** | **QRENLEKEVAEAKHNNETL** | **4.4** |
| **207** | **109** | **QKENLEKEVAEAKHKNETL** | **4.4** |
| **18** | **108** | **QKENLEKEVAEAKHKNETL** | **4.4** |
| **55** | **270** | **EIQEKEAEKDRQQHMYEAF** | **4.3** |
| **222** | **262** | **EIQEKEAEKDRQQHMYEAF** | **4.3** |
| **164** | **88** | **EQQERQKKLEQLEHKYQVE** | **4.2** |
| **37** | **134** | **EVQEKEYNNDGLRHKNDDL** | **3.9** |
| **170** | **91** | **EQKERQKKLEQLEHKYQVE** | **3.9** |
| **M22_enn342** | **132** | **EQKERQKKLEQLEHKYQVE** | **3.9** |
| **M48_enn340** | **132** | **EQKERQKKLEQLEHKYQVE** | **3.9** |
| **M75_enn334** | **132** | **EQKERQKKLEQLEHKYQVE** | **3.9** |
| **M75_enn335** | **132** | **EQKERQKKLEQLEHKYQVE** | **3.9** |
| **M81_enn319** | **129** | **EQKERQKKLEQLEHKYQVE** | **3.9** |
| **M81_enn320** | **129** | **EQKERQKKLEQLEHKYQVE** | **3.9** |
| 79 | 80 | DYSQIEEKLEQFGHDYDKL | 3.6 |
| 87 | 81 | DYSEIEGKLEQFWHDYDKL | 3.6 |
| 105 | 59 | RADKLETENHGLKFQNEKL | 3.4 |
| 105 | 143 | QRVRLEKQVQEKEHNNKTL | 3.4 |
| 207 | 60 | RFEASDLENHKLKFDNDKL | 3.4 |
| 79 | 108 | QRVKLEKQVQEKEHNNKTL | 3.4 |
| 218 | 122 | QRVKLEKQVQEKEHNNKTL | 3.4 |
| 100 | 59 | KADKYEVRNHELEHNNEKL | 3.4 |
| 209 | 80 | DYSQIQEELEQFGHDYDKL | 3.3 |
| 31 | 230 | ENAKKDFELAALGHQLADK | 3.25 |
| 229 | 224 | ENAKKDFELAALGHQLADK | 3.25 |
| 12 | 222 | ENAKKDFELAALGHQLADK | 3.25 |
| 228 | 208 | ENAKKDFELAALGHQLADK | 3.25 |
| 39 | 207 | ENAKKDFELAALGHQLADK | 3.25 |
| 193 | 200 | ENAKKDFELAALGHQLADK | 3.25 |
| M11_enn344 | 132 | EHKERQEKLEQLEHKYQVE | 3.2 |
| M63_enn346 | 132 | EQKERQEKLEQLEHKYQVE | 3.2 |
| M63_enn327 | 129 | EQKERQEKLEQLEHKYQVE | 3.2 |
| M63_enn324 | 129 | EQKERQEKLEQLEHKYQVE | 3.2 |
| M63_enn325 | 129 | EQKERQEKLEQLEHKYQVE | 3.2 |
| M85_enn329 | 129 | EQKERQEKLEQLEHKYQVE | 3.2 |
| M85_enn330 | 129 | EQKERQEKLEQLEHKYQVE | 3.2 |

| | | | |
|---|---|---|---|
| MB5_mrp70 | 61 | EEVIANMSLDKLQHTLAGS | 3.1 |
| MB5_mrp71 | 61 | EEVIANMSLDKLQHTLAGS | 3.1 |
| MB5_mrp72.0 | 61 | EEVIANMSLDKLQHTLAGS | 3.1 |
| MB5_mrp79 | 61 | EEVITNMSLEELQHTLAGS | 3.1 |
| 39 | 57 | EYHRLDTENHTLKHDKEKL | 2.9 |
| 14 | 57 | RAQDLEAKKHALEHQNTKL | 2.9 |
| 14 | 57 | RAQDLEAKNHGLEHQNTKL | 2.9 |
| 63 | 44 | EAQNNNSGKLTLEHKYNAL | 2.8 |
| MA2_mrp160 | 185 | EAETLENLLGSAKHELTEL | 2.7 |
| MA2_mrp161 | 185 | EAETLENLLGSAKHELTEL | 2.7 |
| 26 | 122 | NNKTLQTQNEDLTHENGQL | 2.6 |
| 36 | 104 | ELTEQNKELKAEEHRLITE | 2.6 |
| 57 | 213 | KLGQLNIDNIDLKHELEQE | 2.6 |
| 57 | 185 | ENQDLEEKLKDKEFYLGET | 2.5 |
| MB5_mrp55 | 56 | REKALEEVIAKMPFEELQH | 2.5 |
| MB5_mrp56 | 56 | REKALEEVIAKMPFEELQH | 2.5 |
| MB5_mrp57 | 56 | REKALEEVIAKMPFEELQH | 2.5 |
| MB5_mrp58 | 56 | REKALEEVIAKMPFEELQH | 2.5 |
| MB5_mrp62 | 56 | REKALEEVIAKMPFEELQH | 2.5 |
| MA1_mrp282 | 189 | EAATLENLVGSAKHELTDL | 2.45 |
| MA1_mrp284 | 189 | EAATLENLVGSAKHELTDL | 2.45 |
| MA1_mrp287 | 189 | EAATLENLLGSAKHELTEL | 2.45 |
| MA2_mrp115.0 | 185 | EAATLENLLGSAKHELTDL | 2.45 |
| MA2_mrp117 | 185 | EAATLENLLGSAKHELTDL | 2.45 |
| MA2_mrp118 | 185 | EAATLENLLGSAKHELTDL | 2.45 |
| MA2_mrp119 | 185 | EAATLENLLGSAKHELTDL | 2.45 |
| MA2_mrp136 | 185 | EAATLENLLGSAKHELTDL | 2.45 |
| MA2_mrp139 | 185 | EAATLENLLGSAKHELTDL | 2.45 |
| MA2_mrp140 | 185 | EAATLENLLGSAKHELTDL | 2.45 |
| MA2_mrp141 | 185 | EAATLENLLGSAKHELTGL | 2.45 |
| MA2_mrp143 | 185 | EAATLENLLGSAKHELTGL | 2.45 |
| MA2_mrp144 | 185 | EAATLENLLGSAKHELTGL | 2.45 |
| MA2_mrp145 | 185 | EAATLENLLGSAKHELTGL | 2.45 |
| MA2_mrp155 | 185 | EAATLENLLGSAKHELTEL | 2.45 |
| MA2_mrp156 | 185 | EAATLENLLGSAKHELTEL | 2.45 |
| MA2_mrp159 | 185 | EAATLENLLGSAKHELTEL | 2.45 |
| MA2_mrp167 | 185 | EAATLENLLGSAKHELTDL | 2.45 |
| MA2_mrp168 | 185 | EAATLENLLGSAKHELTDL | 2.45 |
| MA2_mrp171 | 185 | EAATLENLLGSAKHELTDL | 2.45 |
| MA2_mrp172 | 185 | EAATLENLLGSAKHELTEL | 2.45 |
| MA2_mrp174 | 185 | EAATLENLLGSAKHELTDL | 2.45 |
| MA3_mrp293 | 220 | EAATLENLLGSAKHELTDL | 2.45 |
| MA3_mrp295 | 220 | EAATLENLLGSAKHELTDL | 2.45 |
| MA3_mrp296 | 220 | EAATLENLLGSAKHELTDL | 2.45 |
| MA3_mrp297 | 220 | EAATLENLLGSAKHELTEL | 2.45 |
| MA3_mrp298 | 220 | EAATLENLLGSAKHELTEL | 2.45 |
| MB4_mrp6 | 226 | EAATLENLLGSAKHELTEL | 2.45 |
| MB4_mrp7 | 226 | EAATLENLLGSAKHELTEL | 2.45 |
| MB4_mrp8.0 | 226 | EAATLENLLGSAKHELTEL | 2.45 |
| MB4_mrp16 | 226 | EAATLENLLGSAKHELTDL | 2.45 |
| MB4_mrp19 | 226 | EAATLENLLGSAKHELTDL | 2.45 |
| MB4_mrp20.0 | 226 | EAATLENLLGSAKHELTDL | 2.45 |
| MB4_mrp22 | 226 | EAATLENLLGSAKHELTDL | 2.45 |
| MB4_mrp23 | 226 | EAATLENLLGSAKHELTDL | 2.45 |
| MB4_mrp25 | 226 | EAATLENLLGSAKHELTDL | 2.45 |

| | | | |
|---|---|---|---|
| MB4_mrp26 | 226 | EAATLENLLGSAKHELTDL | 2.45 |
| MB4_mrp27 | 226 | EAATLENLLGSAKHELTDL | 2.45 |
| MB4_mrp30 | 226 | EAATLENLLGSAKHELTDL | 2.45 |
| MB4_mrp31.1 | 226 | EAATLENLLGSAKHELTDL | 2.45 |
| MB4_mrp34 | 226 | EAATLENLLGSAKHELTDL | 2.45 |
| MB4_mrp36 | 226 | EAATLENLLGSAKHELTDL | 2.45 |
| MB4_mrp37.1 | 226 | EAATLENLLGSAKHELTDL | 2.45 |
| MB4_mrp39 | 226 | EAATLENLLGSAKHELTDL | 2.45 |
| MB4_mrp40 | 226 | EAATLENLLGSAKHELTEL | 2.45 |
| MB4_mrp41 | 226 | EAATLENLLGSAKHELTDL | 2.45 |
| MB4_mrp42 | 226 | EAATLENLLGSAKHELTDL | 2.45 |
| MB4_mrp43.0 | 225 | EAATLENLLGSAKHELTEL | 2.45 |
| MB4_mrp45.0 | 225 | EAATLENLLGSAKHELTEL | 2.45 |
| MB4_mrp46 | 225 | EAATLENLLGSAKHELTEL | 2.45 |
| MB4_mrp48 | 225 | EAATLENLLGSAKHELTEL | 2.45 |
| MB4_mrp49 | 225 | EAATLENLLGSAKHELTEL | 2.45 |
| MB4_mrp50 | 225 | EAATLENLLGSAKHELTDL | 2.45 |
| MB4_mrp53 | 225 | EAATLENLLGSAKHELTDL | 2.45 |
| MB5_mrp88 | 216 | EAATLENLLGSAKHELTEL | 2.45 |
| MB5_mrp89.1 | 216 | EAATLENLLGSAKHELTEL | 2.45 |
| MB5_mrp91 | 216 | EAATLENLLGSAKHELTEL | 2.45 |
| MB5_mrp93 | 216 | EAATLENLLGSAKHELTDL | 2.45 |
| MB5_mrp94 | 216 | EAATLENLLGSAKHELTDL | 2.45 |
| MB5_mrp96 | 216 | EAATLENLLGSAKHELTDL | 2.45 |
| MB5_mrp98 | 216 | EAATLENLLGSAKHELTEL | 2.45 |
| MB5_mrp99 | 216 | EAATLENLLGSAKHELTEL | 2.45 |
| MB5_mrp100 | 216 | EAATLENLLGSAKHELTEL | 2.45 |
| MB5_mrp101 | 216 | EAATLENLLGSAKHELTEL | 2.45 |
| MB5_mrp102 | 216 | EAATLENLLGSAKHELTEL | 2.45 |
| MB5_mrp103 | 216 | EAATLENLLGSAKHELTEL | 2.45 |
| MB5_mrp55 | 216 | EAATLENLLGSAKHELTEL | 2.45 |
| MB5_mrp56 | 216 | EAATLENLLGSAKHELTEL | 2.45 |
| MB5_mrp57 | 216 | EAATLENLLGSAKHELTEL | 2.45 |
| MB5_mrp58 | 216 | EAATLENLLGSAKHELTDL | 2.45 |
| MB5_mrp59 | 216 | EAATLENLLGSAKHELTEL | 2.45 |
| MB5_mrp61 | 216 | EAATLENLLGSAKHELTDL | 2.45 |
| MB5_mrp63 | 216 | EAATLENLLGSAKHELTDL | 2.45 |
| MB5_mrp66 | 216 | EAATLENLLGSAKHELTDL | 2.45 |
| MB5_mrp68 | 216 | EAATLENLLGSAKHELTEL | 2.45 |
| MB5_mrp69 | 216 | EAATLENLLGSAKHELTEL | 2.45 |
| MB5_mrp70 | 216 | EVATLENLLGSAKHELTEL | 2.45 |
| MB5_mrp71 | 216 | EAATLENLLGSAKHELTEL | 2.45 |
| MB5_mrp72.0 | 216 | EAATLENLLGSAKHELTEL | 2.45 |
| MB5_mrp75 | 216 | EAATLENLLGSAKHELTDL | 2.45 |
| MB5_mrp76 | 216 | EAATLENLLGSAKHELTDL | 2.45 |
| MB5_mrp77 | 216 | EAATLENLLGSAKHELTDL | 2.45 |
| MB5_mrp79 | 216 | EAATLENLLGSAKHELTEL | 2.45 |
| MB5_mrp80 | 216 | EAATLENLLGSAKHELTEL | 2.45 |
| MB5_mrp82 | 216 | EAATLENLLGSAKHELTDL | 2.45 |
| MB5_mrp86.0 | 216 | EAATLENLLGSAKHELTEL | 2.45 |
| 93 | 60 | RLDEQNHKLVNDNHKLVND | 2.4 |

**Table S3. M6 FH-binding pattern in M, Enn and Mrp proteins.**

Red horizontal line indicates cut-off.

| M | Position | Sequence | Score |
|---|---|---|---|
| **6** | **123** | **NKELKAEENRLTTENKGLTKKLSEAEEEAANKERE** | **6.4** |
| **36** | **109** | **NKELKAEEHRLITENRGLTKKLSEAEEESVNKERE** | **6.4** |
| **213** | **95** | **NEDLTREYRRLTQDNRGLTQKNEDLTQKNHRLTQE** | **6.4** |
| **71** | **95** | **NEDLTQEKQRLTSENRGLTKENEDLTQKNHRLTQD** | **6.4** |
| **32** | **67** | **NHQLTQENEKLTQENEKLTKDKEELTQENEKLTKD** | **6.1** |
| **183** | **56** | **YSQLHDDYDKLQEQNGEYLKKIGELEERQKNLEKL** | **5.8** |
| **238** | **57** | **ANNTTVQNIRLRNENKNLKAKNEDLEARLENAMNV** | **5.8** |
| **17** | **88** | **NEELGQEKEKLGQENEELKQEKEKLKTQAAELEET** | **5.7** |
| **17** | **123** | **NREYGAEKDRLVLENRDLENKNRDLENKNRDLEGG** | **5.7** |
| **26** | **144** | **NKDYEAENGRLESDNRRLEGKNKDLEGKNKDLEGK** | **5.7** |
| **115** | **116** | **NEDLTREYDRLTQENRGLTQDKDELSKQKETLGLA** | **5.7** |
| **30** | **165** | **KEDLTREYRRLTQDNRGLTKDREDLTQKNHELSGQ** | **5.7** |
| **197** | **193** | **KEDLTREYRRLTQDNRGLTKDREDLTQKNHELSGQ** | **5.7** |
| M18_enn300 | 96 | KWNLNDEYNKLLDENEKLKEEIGGYLDKQEQLEQL | 5.4 |
| M64_enn306 | 96 | KWNLNDEYNKLLDENEKLKEEIGGYLDKQEQLEQL | 5.4 |
| M80_enn310 | 96 | KWNLNDEYNKLLDENEKLKEEIGGYLDKQEQLEQL | 5.4 |
| M80_enn311 | 96 | KWNLNDEYNKLLDENEKLKEEIGGYLDKQEQLEQL | 5.4 |
| M98_enn314 | 96 | QWNLTEEYNKLHEENERLKEEIGGYLDKQDQLEQL | 5.4 |
| M101_enn300 | 96 | KWNLNDEYNKLLDENEKLKEEIGGYLDKQEQLEQL | 5.4 |
| M123_enn300.1 | 96 | KWNLNDEYNKLLDENEKLKEEIGGYLDKQEQLEQL | 5.4 |
| 158 | 67 | RDNLLGENGKLWDENETLREKQEELEKENEKLDSQ | 5.4 |
| 205 | 55 | KWNLNDEYNKLLDENEKLKEEIGGYLDKQEQLEQL | 5.4 |
| 117 | 59 | YNELSGEYNKLLDQNGNLLDENEILKEKLDKDQEE | 5.4 |
| 100 | 67 | NHELEHNNEKLKTENSDLKTENSKLTSEKEELTQE | 5.4 |
| M54_enn260 | 87 | QFDWEKEYKKLDEDNAKLVEVVEATSLENEKLKSE | 5.2 |
| M71_enn263 | 87 | QFDWEKEYKKLDGDNAKLVEVVEATSLENEKLKSE | 5.2 |
| M71_enn262 | 87 | QFDWEKEYKKLDGDNAKLVEVVEATSLENEKLKSE | 5.2 |
| MA1_mrp247 | 113 | LNNKNEQIAKLTNENAQLKEAVEGYVQTIQNASRE | 5.1 |
| MA1_mrp248 | 113 | LNNKNEQIAKLTNENAQLKEAVEGYVQTIQNASRE | 5.1 |
| MA1_mrp278 | 113 | LNNKNEQIAKLTNENAQLKEAVEGYVQTIQNASRE | 5.1 |
| MA1_mrp279 | 113 | LNNKNEQIAKLTNENAQLKEAVEGYVQTIQNASRE | 5.1 |
| 166 | 48 | QVDWEKEYKKLDEDNAKLVEVVETTSLENEKLKSE | 5 |
| 47 | 67 | IHQLKDDKEKLQSQNENLQSQNENLQSQNENLQSQ | 4.9 |
| 79 | 88 | LEQFGHDYDKLEKENKEYASQLGKNQEEREKLELE | 4.7 |
| 209 | 88 | LEQFGHDYDKLEKENKEYASQLGKNQEEREKLELG | 4.7 |
| 87 | 89 | LEQFWHDYDKLEKENKEYASQLGKNQEEREKLELE | 4.7 |
| 103 | 92 | LEQFGRDYDKLEKENKEYASQLGKNQEEREKLELE | 4.7 |
| 157 | 51 | REQLRQEHDRLEAENSKLLNQTEKLQKKITDLTTE | 4.7 |
| 147 | 51 | KEQLRQEHDRLEIENHKLLNETEKLQKKITDLNTK | 4.7 |
